# SUMOylation regulates Lem2 function in centromere clustering and silencing

**DOI:** 10.1101/2022.11.02.514898

**Authors:** Joanna Strachan, Orsolya Leidecker, Christos Spanos, Clementine Le Coz, Elliott Chapman, Ana Arsenijevic, Haidao Zhang, Ning Zhao, Elizabeth H. Bayne

**Affiliations:** Institute of Cell Biology, School of Biological Sciences, University of Edinburgh, Edinburgh, United Kingdom; Max Planck Institute for Biology of Ageing, Joseph-Stelzmann-Strasse 9b, Cologne 50931, Germany; Wellcome Centre for Cell Biology, Institute of Cell Biology, School of Biological Sciences, University of Edinburgh, Edinburgh, United Kingdom

## Abstract

Regulation by the small modifier SUMO is heavily dependent on spatial control of enzymes that mediate the attachment and removal of SUMO on substrate proteins. Here we show that in fission yeast, delocalisation of the SUMO protease Ulp1 from the nuclear envelope results in centromeric defects that can be attributed to hyper-SUMOylation at the nuclear periphery. Unexpectedly, we find that while this localised hyper-SUMOylation impairs centromeric silencing, it can also enhance centromere clustering. Moreover, both effects are at least partially dependent on SUMOylation of the inner nuclear membrane protein Lem2. Lem2 has previously been implicated in diverse biological processes, including the promotion of both centromere clustering and silencing, but how these distinct activities are coordinated was unclear; our observations suggest a model whereby SUMOylation may serve as a regulatory switch, modulating Lem2 interactions with competing partner proteins to balance its roles in alternative pathways. Our findings also reveal a previously unappreciated role for SUMOylation in promoting centromere clustering.

## Introduction

Uniform segregation of genetic material between daughter cells is essential for cellular proliferation and survival. Centromeres play a critical role in this process by directing the formation of kinetochore complexes, onto which spindles from opposite poles of dividing cells can attach to separate duplicated chromosomes. Centromere-associated heterochromatin contributes to centromere function and further promotes genome stability by silencing repetitive genetic elements and suppressing recombination (Allshire and Madhani, 2018). In addition, increasing evidence is emerging that the physical organisation of centromeres within the cell is also important, with clustering of centromeres during interphase seen in many organisms, often in the vicinity of the nuclear periphery (Fransz et al., 2002; Weierich et al., 2003; Solovei et al., 2004; Kozubowski et al., 2013; Padeken et al., 2013). Although the mechanisms and functions of centromere clustering are yet to be fully elucidated, there is evidence that this spatial organisation can facilitate loading of centromeric proteins (Wu et al., 2022), enhance transcriptional silencing (Padeken et al., 2013), and promote genome stability through prevention of micronuclei formation (Jagannathan et al., 2018).

The fission yeast *Schizosaccharomyces pombe* has proven a powerful model for the study of nuclear organisation. During interphase, *S. pombe* chromosomes adopt the so-called Rabl conformation, whereby all three centromeres are clustered together and anchored at the nuclear periphery adjacent to the spindle pole body (SPB; the microtubule organisation centre in yeast) (Funabiki et al., 1993; Mizuguchi et al., 2015). A key factor in this clustering is Csi1, which appears to provide a physical link between kinetochore and SPB-associated proteins. Deletion of Csi1 results in severe centromere clustering defects as well as defects in chromosome segregation during mitosis (Hou et al., 2012). In addition, the inner nuclear membrane protein Lem2 appears to function in parallel with Csi1 to help cluster centromeres, with cells lacking both Csi1 and Lem2 showing synthetic clustering defects (Barrales et al., 2016). Lem2 is localised throughout the nuclear envelope, but shows Csi1-dependent enrichment at the SPB (Ebrahimi et al., 2018). Lem2 shares similarity with Lap2/Emerin/Man1 (LEM) sub-family of animal cell lamina-associated proteins, and although *S. pombe* lacks nuclear lamina, Lem2 shares conserved functions of lamin-related proteins including maintenance of nuclear envelope structure, peripheral tethering of chromatin, and chromatin silencing (Hiraoka et al., 2011; Gonzalez et al., 2012; Banday et al., 2016; Barrales et al., 2016; Tange et al., 2016; Hirano et al., 2018).

Similar to multicellular eukaryotes, centromeres in *S. pombe* comprise two distinct types of chromatin: a core domain enriched for the centromeric histone variant CENP-A^Cnp1^, and flanking regions of repressive heterochromatin associated with pericentromeric repeats (comprising inner most repeat (*imr*) and outer repeat (*otr*) sequences). A complex network of redundant mechanisms contribute to pericentromeric heterochromatin formation and silencing, including both RNA interference (RNAi) dependent and independent pathways (Reyes-Turcu and Grewal, 2012; Marina et al., 2013; Allshire and Ekwall, 2015; Chalamcharla et al., 2015; Martienssen and Moazed, 2015; Tucker et al., 2016; Taglini et al., 2020). Lem2 has also been implicated in pericentromeric silencing, with deletion of Lem2 causing loss of silencing most prominently in the *imr* region (Banday et al., 2016; Barrales et al., 2016). Interestingly, this role of Lem2 in silencing appears to be independent of its role in tethering centromeres at the nuclear periphery: while the N-terminal HeH/LEM domain of Lem2 associates with centromeric chromatin and is required for chromatin tethering (Barrales et al., 2016; Tange et al., 2016), the C-terminal Man1-Src1p-C-terminal (MSC) domain of Lem2 is sufficient to mediate centromere silencing, possibly *via* recruitment of repressive chromatin factors (Banday et al., 2016; Barrales et al., 2016). A growing number of partner proteins have been implicated in contributing to the diverse functions of Lem2; for example, binding to the inner nuclear membrane protein Bqt4 helps mobilise Lem2 around the nuclear envelope and mediates telomere-related functions (Ebrahimi et al., 2018; Hirano et al., 2018), whilst interaction of Lem2 with the RNA surveillance factor Red1 was recently found to contribute to regulation of meiotic transcripts (Martin Caballero et al., 2022). However, how Lem2 is differentially targeted for roles in these varied processes is yet to be fully elucidated.

Lem2 is one of many centromere and kinetochore associated proteins found to be subject to SUMOylation in fission yeast (Kohler et al., 2015). SUMO is a small protein modifier that is similar in structure to ubiquitin, and like ubiquitin, can be covalently attached to lysine residues in substrate proteins through a cascade of E1-activating, E2-conjugating and E3 ligase enzymes. However, whereas a relatively large and diverse array of E3 ligases confer substrate specificity for ubiquitination, the complement of E3 SUMO ligases is much smaller, reflecting a greater role for spatial control in substrate definition (Psakhye and Jentsch, 2012; Jentsch and Psakhye, 2013). Whilst a classic fate of ubiquitinated proteins is proteasome-mediated degradation, SUMOylation is more often associated with modulating protein-protein interactions, either negatively, for example by obscuring a binding surface (Pichler et al., 2005), or more commonly positively *via* non-covalent interaction of SUMO with a SUMO interacting motif (SIM). Individual SUMO-SIM interactions often act synergistically to enhance binding affinities (Hecker et al., 2006; Yau et al., 2021), and these interactions can also influence protein localisation (Mahajan et al., 1998; Matunis et al., 1998). In addition, cross-talk between SUMO and ubiquitin modifications can occur; for example, in some circumstances polySUMOylated proteins can be recognised by SUMO-targeted ubiquitin ligases (STUbLs), with subsequent ubiquitination driving extraction or degradation of target proteins (Uzunova et al., 2007; Perry et al., 2008; Nie et al., 2012; Kohler et al., 2013; Kohler et al., 2015; Nie and Boddy, 2015).

Like other post-translational modifications, SUMOylation is dynamic and reversible. SUMO deconjugation is performed by the conserved ULP/SENP family of SUMO-specific proteases, which includes six SENP family proteins in human, and Ulp1 and Ulp2 in yeast. Differences in substrate specificity between these enzymes often appear to be determined by their distinct subcellular localisations (Li and Hochstrasser, 2003; Hickey et al., 2012). In *S. cerevisiae*, Ulp2 is localised throughout the nucleoplasm, but shows highest activity towards SUMO-SUMO linkages and therefore the shortening of polySUMO chains. In contrast, Ulp1 shows broad specificity, removing SUMO from substrate proteins as well as processing SUMO precursors; however, it is spatially restricted, being localised primarily to the inner surface of the nuclear pore complex. Loss of this localisation results in a major shift in substrates, indicating that the physical location of Ulp1 normally restricts its activity towards certain SUMOylated proteins whilst enabling cleavage of others (Li and Hochstrasser, 2003).

Centromere and kinetochore associated factors are reported to be enriched amongst SUMOylated proteins in several species (Azuma et al., 2003; Montpetit et al., 2006; Zhang et al., 2008; Mukhopadhyay et al., 2010; Ban et al., 2011; Wan et al., 2012; Li et al., 2016; Restuccia et al., 2016), and both SUMO E3 ligase and SUMO protease enzymes have been found to co-localise with kinetochores (Joseph et al., 2004; Agostinho et al., 2008; Ban et al., 2011; Cubenas-Potts et al., 2013; Suhandynata et al., 2019). Indeed, the budding yeast genes encoding both SUMO and Ulp2 were first identified as high copy suppressors of a temperature sensitive mutation in Mif2 (Meluh and Koshland, 1995), the ortholog of mammalian centromere protein CENP-C. It was subsequently shown that the human ortholog of Ulp2, SENP6, is required for proper centromere assembly, by preventing hyper-SUMOylation of multiple proteins within the constitutive centromere associated network (CCAN) to enable their assembly at centromeres (Liebelt et al., 2019). Conversely, recent evidence suggests that SUMOylation of kinetochore protein Nuf2 is required to promote recruitment of SIM-domain containing centromere-associated protein CENP-E, essential for proper alignment of chromosomes in metaphase (Subramonian et al., 2021). Hence the role of SUMO at centromeres appears multifaceted and is yet to be fully defined.

In *S. pombe* there is a single SUMO isoform (Pmt3), and the majority of SUMOylation is directed by one E3 SUMO ligase, Pli1. Deletion of Pli1 is associated with centromere-related defects including impaired silencing at pericentromeric (*imr*) regions and sensitivity to the microtubule destabilising drug thiabendazole (TBZ) (Xhemalce et al., 2004). Similar defects are seen in cells lacking either the SUMO protease Ulp1 (Han et al., 2010), or the nucleoporin Nup132 that is required to tether Ulp1 to the nuclear periphery (Han et al., 2010; Nie and Boddy, 2015). It has previously been proposed that depletion of Pli1, and hence a reduction in SUMOylation, accounts for the defects in all three backgrounds, since it was shown that Nup132-tethered Ulp1 functions to antagonise Pli1 auto-SUMOylation, and in *nup132Δ* cells polySUMOylated Pli1 accumulates and is subject to STUbL-mediated degradation (Nie and Boddy, 2015). However, the release of Ulp1 from the nuclear periphery in *nup132Δ* cells results not only in a reduction in global SUMOylation (linked primarily to destabilisation of Pli1), but also an increase in accumulation of SUMOylated proteins at the nuclear periphery (Kramarz et al., 2021). Indeed, it was recently demonstrated that impaired processing of stalled replication forks in *nup132Δ* cells can be attributed to loss of deSUMOylation activity at the nuclear periphery, with deletion of Pli1 rescuing the defects (Kramarz et al., 2021). Hence questions remain about precisely how alteration of the SUMO landscape in *nup132Δ* cells impacts on centromere function.

Towards investigating the role of SUMOylation in centromere function in *S. pombe*, here we have directly tested whether centromeric defects in *nup132Δ* cells are a result of either Pli1 stabilisation and hence reduced SUMOylation, or conversely enhanced SUMOylation at the nuclear periphery. We present multiple lines of evidence that localised hyper-SUMOylation is the primary driver of centromeric defects in this background, and identify Lem2 as a key substrate whose SUMOylation contributes to defects in centromere silencing. Unexpectedly, we find that contrary to the detrimental effects on centromeric silencing, hyper-SUMOylation can enhance centromere clustering, helping to rescue clustering defects in cells lacking Csi1. Interestingly, this effect is again at least partially mediated through SUMOylation of Lem2. Our results reveal a previously unappreciated role for SUMOylation in promoting centromere clustering, and suggest that SUMOylation may provide a mechanism for coordination of the diverse functions of Lem2, possibly influencing the balance of Lem2 interactions with alternate partner proteins.

## Results

### Centromere defects in *nup132∆* cells are not explained by destabilisation of Pli1

Defects in centromere silencing in *nup132∆* cells were previously proposed to arise as a result of destabilisation of Pli1, since loss of Nup132-dependent localisation of SUMO protease Ulp1 has been shown to lead to increased Pli1 auto-SUMOylation and hence STUbL-dependent degradation (Nie and Boddy, 2015). In order to test this model, we sought to specifically block the ubiquitin-mediated destabilisation of Pli1.

Although Pli1 has been shown to be subject to ubiquitin-dependent degradation (Nie and Boddy, 2015), the specific ubiquitination sites have not been identified. To address this, and because such an analysis has not previously been reported, we performed global identification of protein ubiquitination sites in fission yeast under physiological conditions, by adapting previously-described strategies for enrichment of ubiquitinated peptides based on affinity purification of peptides bearing the diGly moiety left by trypsin digestion of ubiquitin (Udeshi et al., 2013). Whole-cell protein extracts were prepared under denaturing conditions, and sequentially digested with Lys C and trypsin. The resulting peptides were fractionated and subjected to diGly-IP, from which bound peptides were eluted and analysed by liquid chromatography-tandem mass spectrometry (LC-MS/MS) (Fig S1A). We performed eight independent analyses, identifying a total of 5116 ubiquitination sites on 1801 proteins (Table S1). By comparison, the largest previous study of ubiquitination sites in fission yeast, which employed tagged and over-expressed ubiquitin, identified 1200 ubiquitination sites on 494 proteins (Beckley et al., 2015) (Fig S1B). Hence our approach was both more physiological and more sensitive, resulting in identification of a greater number of ubiquitination sites on a wider range of proteins (representing approximately 35% of the *S. pombe* proteome).

From our global analyses we identified three ubiquitinated residues in Pli1: K15, K169 and K214 (Fig 1A). To test whether ubiquitination of these residues promotes degradation of Pli1 in *nup132∆* cells, we replaced endogenous Pli1 with a mutant version in which the three identified lysine residues were mutated to arginine (Pli1^K3R^). Both wild-type and mutant Pli1 were C-terminally FLAG-tagged. Western blot analysis indicated that in wild-type cells, Pli1^K3R^ mutant protein levels are similar to wild-type Pli1. However, in *nup132∆* cells, whereas wild-type Pli1 is destabilised, Pli1^K3R^ levels remain high (Fig 1B). That mutation of these three residues was sufficient to stabilise Pli1 confirms that these sites are responsible for the ubiquitin-dependent degradation of Pli1 in the *nup132∆* background.

**Figure 1.**
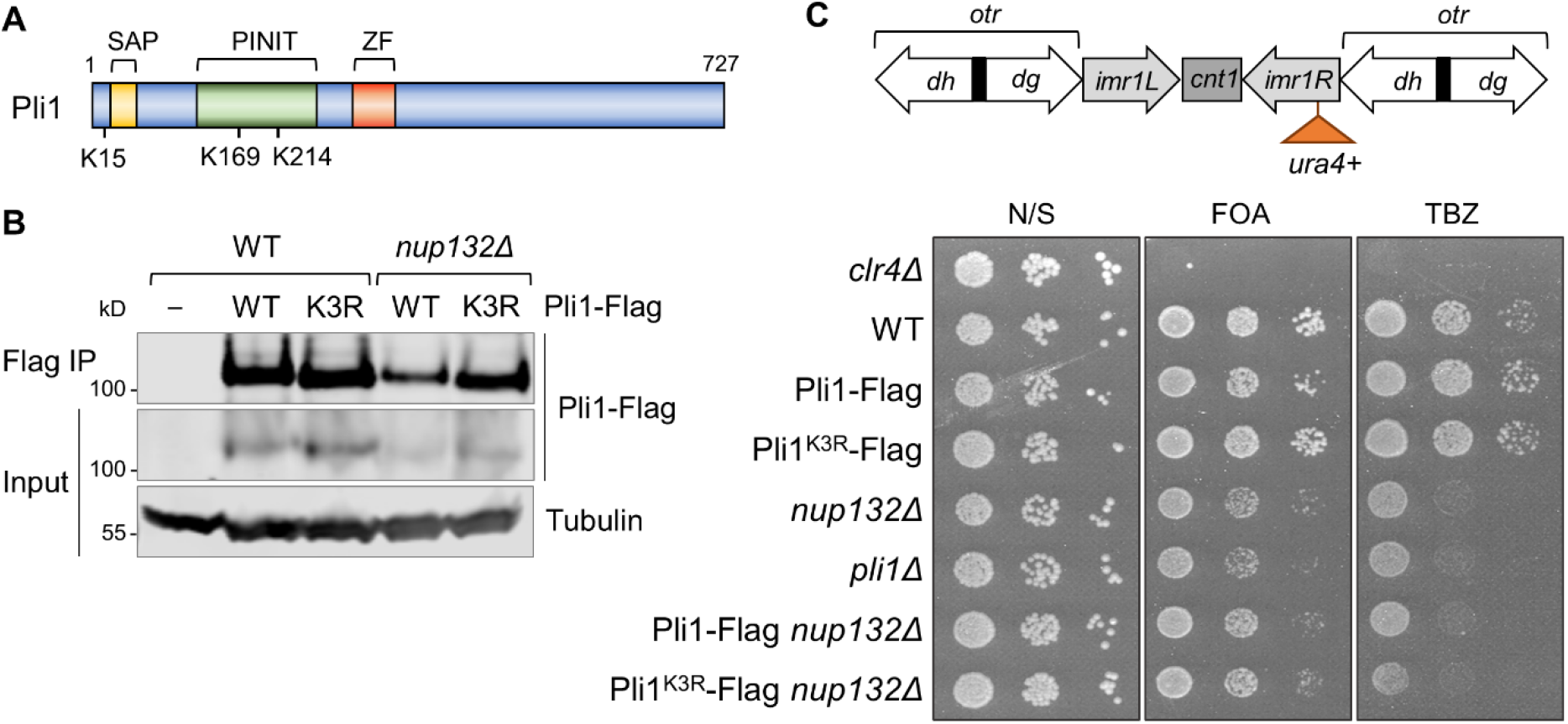
Stabilisation of Pli1 is not sufficient to rescue centromere defects in *nup132∆* cells. **(A)** Schematic indicating the positions of the three ubiquitinated lysine residues within Pli1, determined by proteomic analysis of diGly peptides. Also indicated are the relative positions of the Pli1 SAP domain, PINIT domain and MIZ-type zinc finger (ZF) domain. **(B)** Western blot analysis of levels of Pli1-Flag, or Pli1^K3R^-Flag, immunoprecipitated from wild-type or *nup132∆* cells. Tubulin (anti-Tat1) serves as a loading control. **(C)** Assays for silencing at *imr1:ura4*^*+*^ and TBZ sensitivity. The schematic shows the position of the *imr1:ura4*^*+*^ reporter in centromere 1, relative to outer repeats (*otr; dg and dh*), inner repeats (*imr*), and central core (*cnt*). Loss of silencing results in increased expression of *ura4*^*+*^ and therefore decreased growth in the presence of the counter-selective drug 5-FOA. Plates are non-selective (N/S) or supplemented with either 5-FOA or the microtubule destabilising drug thiabendazole (TBZ).

We next tested whether the stabilising Pli1^K3R^ mutation is also sufficient to rescue the centromere silencing defects observed in *nup132∆* cells. In wild-type cells, a *ura4*^*+*^ reporter gene inserted into the heterochromatic inner-most repeat region of centromere one (*imr:ura4*^*+*^) is repressed, and cells consequently grow well on media containing the counter-selective drug 5-FOA. As reported previously (Xhemalce et al., 2004; Nie and Boddy, 2015), deletion of either *nup132*^*+*^ or *pli1*^*+*^ results in reduced growth in the presence of 5-FOA, indicating increased expression of *ura4*^*+*^ and hence loss of silencing (Fig 1C). If the silencing defect in *nup132∆* cells is due to destabilisation of Pli1, we would expect it to be rescued by expression of the stabilised Pli1^K3R^ mutant. However, this was not the case: *nup132∆* cells expressing Pli1^K3R^-Flag displayed a defect in silencing equivalent to those expressing wild-type Pli1-Flag. In contrast, expression of Pli1^K3R^-Flag in an otherwise wild-type background did not affect silencing. Similarly, both *nup132∆* and *pli1∆* cells show sensitivity to the microtubule stabilising drug TBZ consistent with defects in centromere function, and this TBZ sensitivity is also not rescued by the Pli1^K3R^ mutation (Fig 1C). Therefore contrary to previous assumptions, the centromere defects in *nup132∆* cells do not appear to be due to destabilisation of Pli1.

### Suppression of hyper-SUMOylation in general, or Lem2 SUMOylation in particular, rescues centromere defects in *nup132∆* cells

Since the centromere defects in *nup132∆* cells appeared not to relate to destabilisation of Pli1 and hence reduced global SUMOylation, we reasoned that they might rather relate to increased SUMOylation at the nuclear periphery due to loss of nuclear membrane tethered Ulp1 deSUMOylase activity. This would be consistent with the known association of *S. pombe* centromeres with the nuclear periphery, adjacent to the SPB (Hou et al., 2013; Mizuguchi et al., 2015). To investigate this model, we first tested whether over-expression of Ulp1 is sufficient to rescue the centromere defects in *nup132∆* cells. Interestingly, while Ulp1 over-expression had no effect in wild-type cells, it partially suppressed the TBZ sensitivity of *nup132∆* cells, consistent with the defect resulting from hyper-SUMOylation (Fig 2A). That TBZ sensitivity was not fully suppressed is not entirely unexpected given that Ulp1 is not targeted to the nuclear periphery in this background. We were unable to test whether Ulp1 over-expression suppresses the defect in silencing at *imr:ura4*^*+*^, since this experiment required growth in minimal media but silencing defects in *nup132∆* cells were only observed in rich media (Fig S2). As a further test of the model, we also generated a plasmid expressing a ‘lysine-less’ SUMO (Pmt3^KallR^), in which each of the lysine residues in Pmt3 are replaced by arginine such that SUMO can attach to substrates, but cannot form lysine-linked polySUMO chains. We expressed wild-type and mutant SUMO in “mature” form (terminating in the diGly motif), to avoid any impact of Ulp1 mis-localisation on SUMO maturation. Strikingly, while expression of lysine-less SUMO caused a slight increase in TBZ sensitivity in otherwise wild-type cells, it clearly suppressed the stronger TBZ sensitivity seen in *nup132∆* cells (Fig 2B). Thus these results are consistent with a model in which centromere defects in *nup132∆* cells are primarily due to localised reduction in Ulp1 deSUMOylation activity at the nuclear periphery.

**Figure 2.**
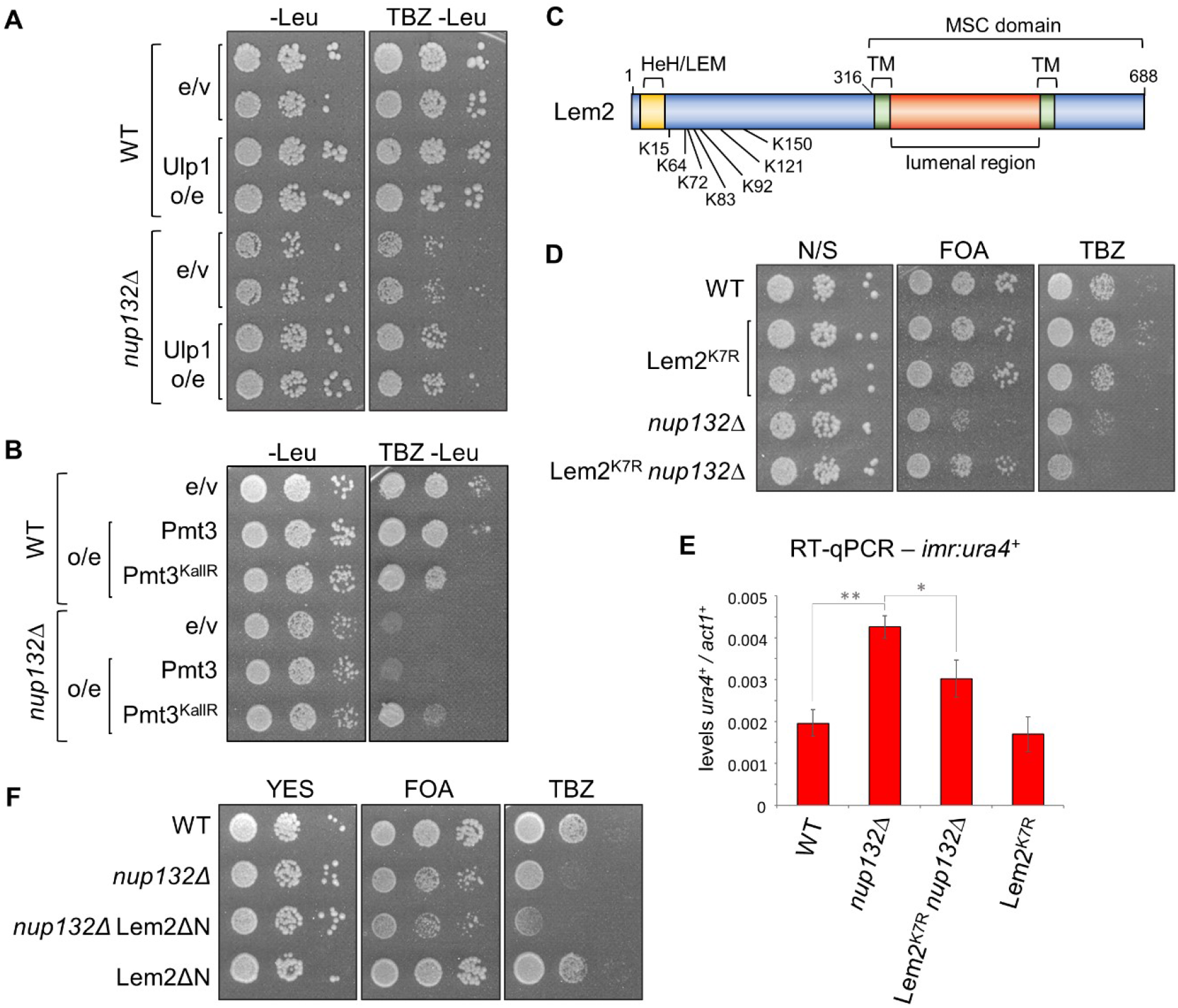
Suppression of hyper-SUMOylation in general, or Lem2 SUMOylation in particular, rescues centromere defects in *nup132∆* cells. **(A)** and **(B)** TBZ sensitivity assay. Wild-type or *nup132∆* cells carrying empty vector (e/v), or over-expressing (o/e) Ulp1, wild-type Pmt3, or Pmt3 in which all lysines have been mutated to arginine (Pmt3^KallR^), were plated on media lacking leucine (-Leu) for maintenance of the plasmid, with or without addition of the microtubule-destabilising drug thiabendazole (TBZ). **(C)** Schematic indicating the positions of the known SUMOylation sites within Lem2. Also indicated are the positions of the HeH/LEM domain and the C-terminal Man1-Src1p-C-terminal (MSC) domain, which includes two transmembrane (TM) domains. **(D)** Assays for silencing at *imr1:ura4*^*+*^ and sensitivity to TBZ. Plates are non-selective (N/S) or supplemented with either 5-FOA or TBZ; loss of silencing results in increased expression of *ura4*^*+*^ and therefore decreased growth in the presence of 5-FOA. **(E)** RT-qPCR analysis of *imr1:ura4*^*+*^ transcript levels, relative to *act1*^*+*^. Data plotted are the mean +/- one standard deviation from three replicates. Asterisks denote *p* ≤ 0.05 (*) and *p* ≤ 0.01 (**) from Student’s *t*-test analysis. **(F)** Assays for silencing at *imr1:ura4*^*+*^ and sensitivity to TBZ. Plates are non-selective (N/S) or supplemented with either 5-FOA or TBZ. Lem2ΔN represents a deletion of the first 307 amino acids of Lem2, leaving only the MSC domain.

We next set out to identify candidate proteins whose hyper-SUMOylation could account for the defects in centromere function in *nup132∆* cells. We searched for candidates meeting three key criteria: (1) a known role in silencing specifically at the centromere *imr* region, similar to Nup132; (2) known localisation to the nuclear periphery; and (3) known to be subject to SUMOylation in *S. pombe*. Interestingly, applying these criteria we identified one clear candidate: the inner nuclear membrane protein Lem2. Lem2 guides nuclear membrane assembly, directly interacts with centromeric and telomeric chromatin to anchor it at the nuclear periphery and crucially plays a role in heterochromatin silencing (Hiraoka et al., 2011; Gonzalez et al., 2012; Banday et al., 2016; Barrales et al., 2016; Tange et al., 2016; Ebrahimi et al., 2018; Hirano et al., 2018; Pieper et al., 2020). Lem2 has been reported to associate with Nup132 (Iglesias et al., 2020), and strikingly, effects of Lem2 deletion on centromere silencing have also been reported to be nutrition-dependent, with stronger defects observed in rich media than in minimal media (Tange et al., 2016). This mirrors our observations for *nup132∆* cells (Fig S2), strongly suggesting that they may function in the same pathway for centromere silencing.

A previous study identified a cluster of seven SUMOylation sites in the N-terminus of Lem2 (Kohler et al., 2015) (Fig 2C). To investigate whether silencing defects in *nup132∆* cells might be a consequence of increased SUMOylation of Lem2 at these sites, we replaced endogenous Lem2 with a mutant version in which each of the seven SUMOylated lysines were replaced with arginine (Lem2^K7R^). The Lem2^K7R^ mutant alone showed no defect in silencing of the *imr:ura4*^*+*^ reporter, as assessed by growth on FOA. Strikingly, when expressed in *nup132∆* cells, Lem2^K7R^ rescued silencing, restoring growth on FOA to near wild-type levels (Fig 2D). This was confirmed by RT-qPCR analysis of *imr:ura4*+ transcript levels, which were elevated in *nup132∆* cells consistent with loss of silencing, but reduced to nearer wild-type levels in *nup132∆* Lem2^K7R^ cells (Fig 2E). These observations suggest that centromere silencing defects in *nup132∆* cells are linked to Lem2 SUMOylation.

Interestingly, while expression of Lem2^K7R^ rescued the centromere silencing defects in *nup132∆* cells, it did not rescue TBZ sensitivity (Fig 2D). Rather, TBZ sensitivity was exacerbated in *nup132∆* Lem2^K7R^ cells, suggesting that other defects, possibly in a pathway functioning in parallel with Lem2, underlie the TBZ sensitivity caused by deletion of Nup132.

Previous studies have shown that while the N-terminus of Lem2 plays the dominant role in tethering chromatin to the nuclear periphery, the C-terminal MSC domain alone is sufficient for Lem2-mediated centromeric silencing (Banday et al., 2016; Barrales et al., 2016). Since Lem2 SUMOylation sites lie in the N-terminal region, we expected that deletion of the Lem2 N-terminal domain including the SUMOylation sites should also be effective in rescuing the silencing defects in *nup132∆* cells. However, interestingly, this was not the case: while deletion of the Lem2 N-terminal domain (Lem2 ΔN; comprising MSC domain only) did not itself impair centromeric silencing, it also did not rescue the silencing defects in *nup132∆* cells (Fig 2F). Thus suppressing Lem2 SUMOylation rescues silencing defects in *nup132∆* cells, but only when the N-terminal domain is present.

The observation that the normally dispensable N-terminal domain of Lem2 is required for centromeric silencing in *nup132∆* cells expressing Lem2^K7R^ argues against the possibility that mutation of Lem2 SUMO sites simply serves to stabilise Lem2 in *nup132Δ* cells, by blocking STUbL-mediated degradation as seen for Pli1. Consistent with this, and in contrast to our observations for Pli1, no difference in protein stability was observed between GFP-tagged wild-type Lem2 and Lem2^K7R^ in *nup132∆* cells (Fig S3A). Additionally, live cell imaging of Lem2-GFP or Lem2^K7R^-GFP revealed no apparent difference in Lem2 localisation in wild-type versus *nup132∆* cells (Fig S3B). We therefore suspect that SUMOylation may rather impact on Lem2 function by affecting protein-protein interactions.

### Increased SUMOylation enhances centromere clustering in *csi1∆* cells

As well as its function in silencing, Lem2 plays an important role in facilitating clustering of centromeres at the SPB (Barrales et al., 2016). Given the Lem2-dependent defect in pericentromeric silencing in *nup132∆* cells, we next tested whether centromere clustering is also affected in this background. Live cell imaging was performed on cells expressing GFP-Cnp1 to visualise centromere positioning, and Sid4-RFP as a marker of the SPB. Consistent with previous reports (Barrales et al., 2016), while 100% of wild-type cells displayed a single SPB-associated GFP-Cnp1 focus (representing the three clustered centromeres) adjacent to the SPB, deletion of *lem2*^*+*^ resulted in a greater proportion of cells with two or even three GFP-Cnp1 foci, indicating centromere clustering defects (Fig 3A and 3B). Interestingly, *nup132∆* cells also exhibited a mild but significant declustering of centromeres. However, whereas the silencing defects in *nup132∆* cells were rescued by expression of Lem2^K7R^, the centromere clustering defects were not (Fig 3B, compare bars 3 and 5). Expression of Lem2^K7R^ alone was also not associated with any centromere clustering defects. Consequently, we find no evidence that the mild centromere clustering defects in *nup132∆* cells are dependent on Lem2 SUMOylation. Interestingly, we also noticed that unlike the silencing defects, the centromere clustering defects in *nup132∆* cells are independent of nutritional status, occurring also in cells grown in minimal media (Fig 3D, bar 5), consistent with there being a different underlying cause.

**Figure 3.**
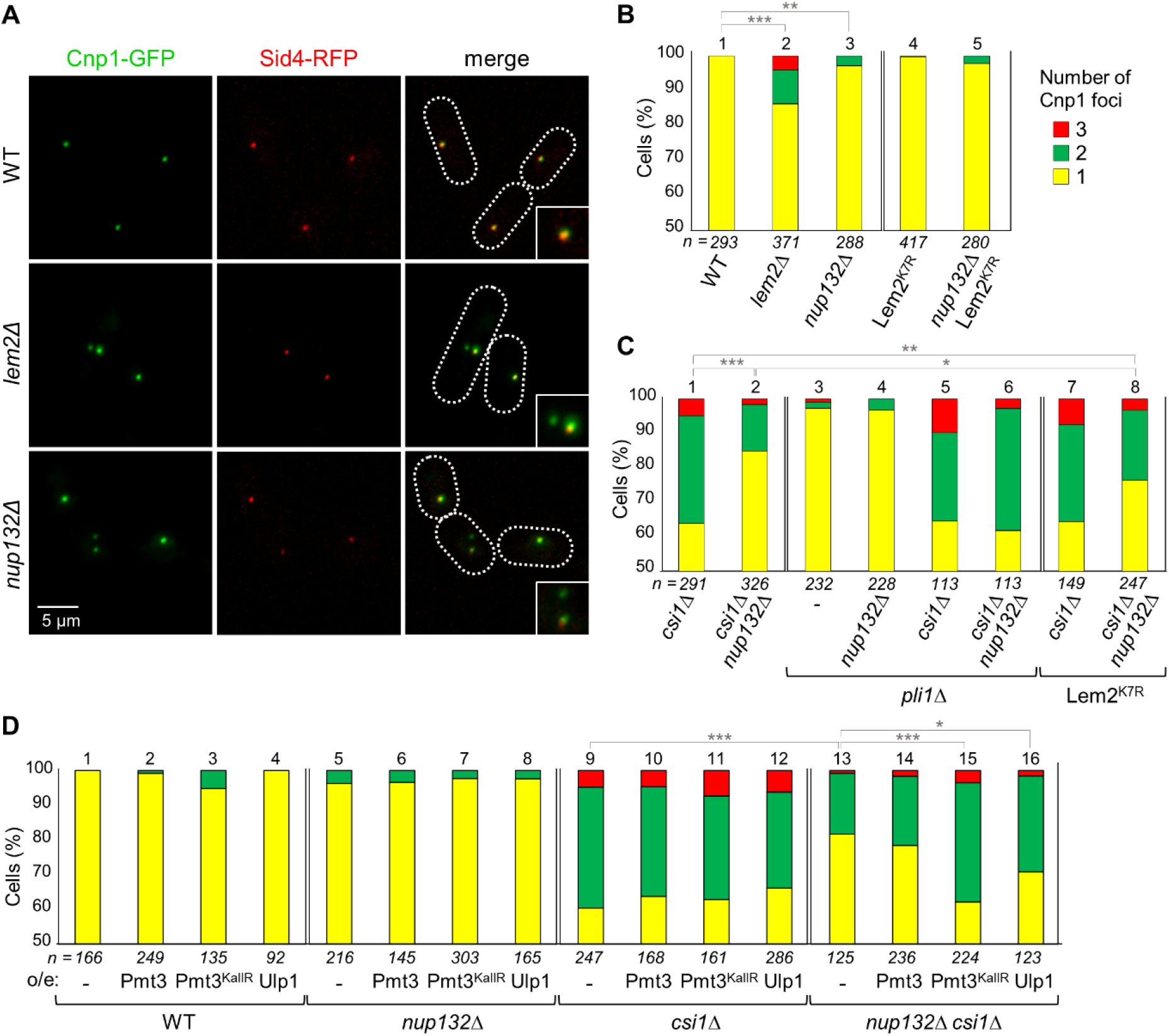
Increased SUMOylation rescues centromere clustering defects in *csi1Δ* cells. **(A)** Representative images from two-colour live cell imaging of GFP-Cnp1 (centromere) and Sid4-RFP (SPB). Dotted lines indicate cell boundaries. **(B) – (D)** Quantification of cells displaying one, two or three Cnp1 foci. Shown are percentages based on analysis of *n* cells. In (D), cells either carry empty vector (-), or over-express Ulp1, wild-type Pmt3, or Pmt3 in which all lysines have been mutated to arginine (Pmt3^KallR^), and are grown in media lacking leucine for maintenance of the plasmid. Asterisks denote *p* ≤ 0.05 (*), *p* ≤ 0.01 (**) and *p* ≤ 0.001 (***) from χ2 test analysis.

It has been shown previously that Lem2 and Csi1 function in parallel pathways to promote centromere clustering (Barrales et al., 2016). Since the clustering defects we observed in *nup132∆* cells appear to be unexpectedly Lem2-independent, we questioned whether *nup132*^*+*^ deletion may instead affect the Csi1-dependent centromere clustering pathway. To test this we generated *nup132∆ csi1∆* double mutant cells, with the expectation that if Nup132 and Csi1 act in parallel pathways, we should observe synthetic defects compared to the single mutants, similar to what has been seen for *lem2∆ csi1∆* cells; and conversely, if Csi1 and Nup132 function in the same pathway, we would observe no further augmentation of clustering defects. Unexpectedly, and contradicting either hypothesis, we found that deletion of *nup132*^*+*^ substantially suppresses centromere declustering defects in *csi1∆* cells, observing normal clustering in 85.0% of *nup132∆ csi1∆* double mutant cells, as compared to only 63.9% of *csi1∆* single mutant cells (Fig 3C).

We next sought to understand the mechanism by which deleting *nup132*^*+*^ rescues centromere clustering defects in *csi1∆* cells. We previously found evidence that the TBZ sensitivity of *nup132∆* cells is related to increased SUMOylation at the nuclear periphery. To test whether hyper-SUMOylation also accounts for the alleviation of centromere clustering defects in *csi1∆* cells upon deletion of *nup132*^*+*^, we tested whether the alleviation is inhibited by manipulations that reduce polySUMO accumulation. Indeed, while over-expression of either Ulp1 (increasing de-SUMOylation), or Pmt3^KallR^ (suppressing polySUMOylation) had little effect on centromere clustering in wild-type or *csi1∆* or *nup132∆* single mutant backgrounds, it was sufficient to fully suppress the rescue of centromere clustering in *csi1∆ nup132∆* double mutant cells (Fig 3D, compare bar 13 to bars 15 and 16). The rescue was also entirely suppressed by deletion of *pli1*^*+*^, repressing global SUMOylation (Fig 3C, compare bars 2 and 6). Together these observations strongly suggest that increased polySUMOylation at the nuclear periphery in *nup132∆* cells can rescue centromere clustering defects caused by loss of Csi1.

One known function of polySUMOylation is the recruitment of STUbLs for extraction and/or degradation of target proteins (Uzunova et al., 2007; Perry et al., 2008; Nie et al., 2012; Kohler et al., 2013; Kohler et al., 2015; Nie and Boddy, 2015). We therefore tested whether the rescue of centromere clustering in *csi1∆* cells upon deletion of *nup132*^*+*^ is dependent on the STUbL Slx8 or on Ufd1 (a component of a ubiquitin–selective chaperone) (Nie et al., 2012; Kohler et al., 2015). However, neither the deletion of *slx8*^*+*^, nor expression of mutant Ufd1 (*ufd1∆Ct*^*213-342*^, lacking the C-terminal domain that mediates interaction with SUMO (Kohler et al., 2013)), prevented the rescue (Fig S4, compare bars 2 and 3, and 5 and 6). Thus the mechanism of rescue does not appear to involve STUbL-dependent protein removal, and rather, SUMO might promote centromere clustering by influencing protein-protein interactions and/or localisation.

### SUMOylation of Lem2 contributes to rescue of centromere clustering in *csi1∆* cells

We previously found that centromere silencing defects in *nup132∆* cells relate to Lem2 SUMOylation, whereas the clustering defects in *nup132∆* cells are not a result of Lem2 SUMOylation. We therefore questioned whether the hyper-SUMOylation of Lem2 in *nup132∆* cells is in fact facilitating the rescue of centromere clustering defects caused by absence of Csi1. Strikingly, we found that while expression of Lem2^K7R^ does not affect clustering in wild-type or *csi1∆* cells, it does partially suppress the rescue seen in *nup132∆ csi1∆* cells, with clustering reduced from 85.0% to 76.5% upon expression of Lem2^K7R^ (Fig 3C, compare bars 1, 2 and 8). This suggests that Lem2 is at least one substrate whose SUMOylation helps to promote centromere clustering in the absence of Csi1.

One function of Csi1 is to help stabilise Lem2 at the SPB (Ebrahimi et al., 2018). Since SUMOylation can influence protein localisation, we next examined whether the rescue of clustering defects in *nup132∆ csi1∆* cells might relate to enhanced localisation of Lem2 at the SPB. Whereas GFP-tagged Lem2 was found to reliably localise to the SPB (indicated by SPB marker Sid4-RFP) in wild-type and *nup132Δ* cells, we observed a loss of Lem2 localisation at the SPB upon deletion of *csi1*^*+*^, consistent with previous findings (Ebrahimi et al., 2018). However, remarkably, while only 75.0% of *csi1∆* cells show normal Lem2 localisation at the SPB, this was increased to 85.2% in *nup132∆ csi1∆* cells (Fig 4A and B). That Lem2 localisation is partially rescued in the *nup132∆* background is consistent with this contributing to the rescue of centromere clustering. To confirm that this enhancement in Lem2 SPB localisation, like the suppression of centromere clustering defects, is dependent on Lem2 SUMOylation, we analysed Lem2^K7R^-GFP subcellular localisation in *nup132∆ csi1∆* cells. Strikingly, we found a reduction in SPB localisation of Lem2^K7R^-GFP as compared to wild-type Lem2-GFP in *nup132∆ csi1∆* cells, closely mirroring the effects on centromere clustering (Fig 4B). Interestingly, we also noted a small reduction in SPB localisation of Lem2^K7R^-GFP as compared to wild-type Lem2-GFP in *csi1∆* cells, consistent with Lem2 SUMOylation contributing to its localisation to SPB in the absence of Csi1. Together, these observations support a model in which hyper-SUMOylation of Lem2, induced by deletion of *nup132*^*+*^, can rescue centromere clustering defects in *csi1∆* cells by increasing the localisation of Lem2 to the SPB and amplifying its contribution to centromere clustering.

**Figure 4.**
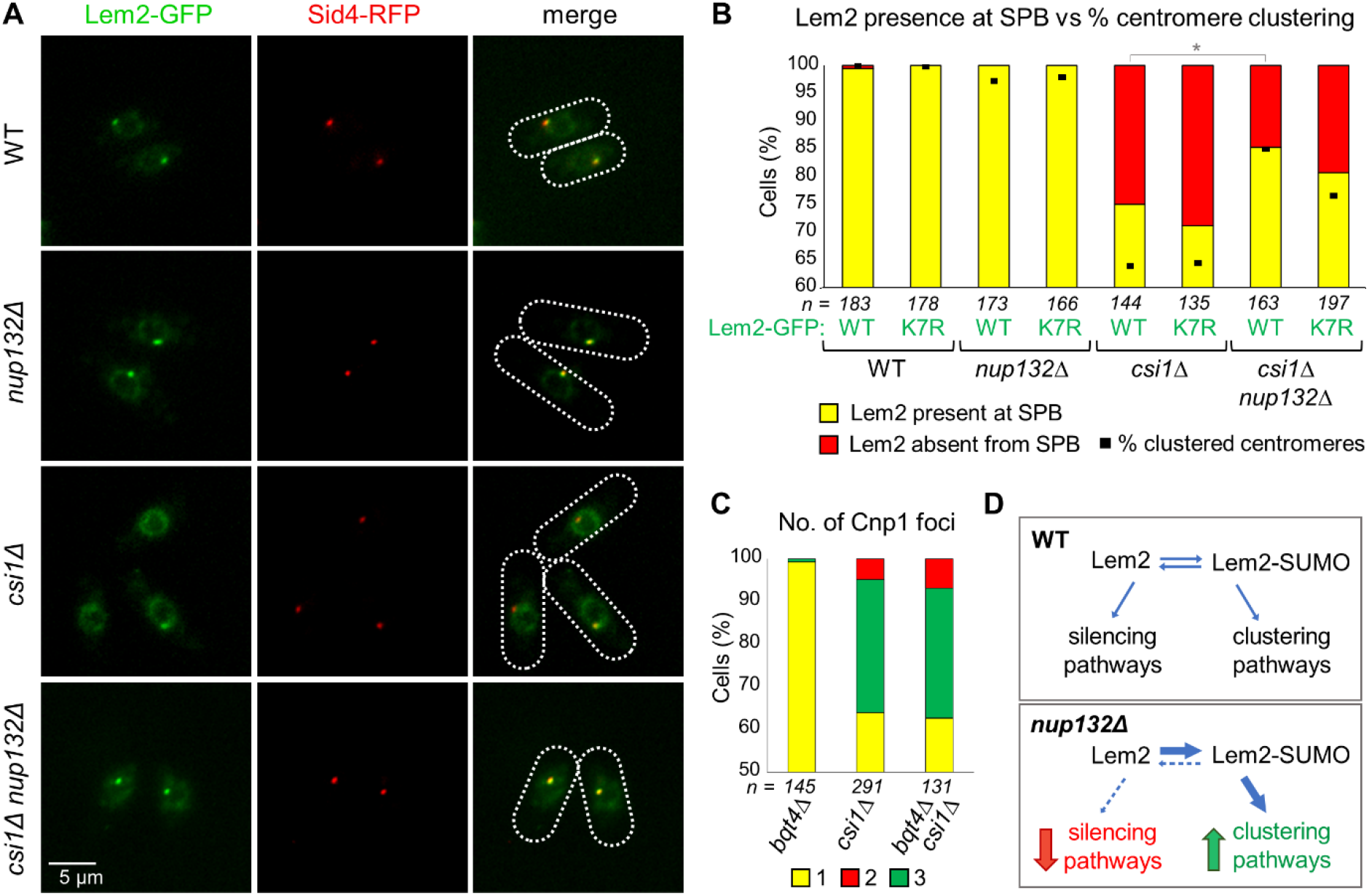
SUMOylation enhances Lem2 localisation at the SPB. **(A)** Representative images from two-colour live cell imaging of Lem2-GFP and Sid4-RFP (SPB marker). Dotted lines indicate cell boundaries. **(B)** Quantification of cells displaying co-localisation of Lem2-GFP and Sid4-RFP. Bars indicate percentages based on analysis of *n* cells. Asterisk (*) denotes *p* ≤ 0.05 from χ2 test analysis. For comparison, black dots indicate percentage of cells in these strains displaying one Cnp1 focus (indicating proper centromere clustering; data are the same as in Fig 3B and C). **(C)** Quantification of cells displaying one, two or three Cnp1 foci, based on live-cell imaging of GFP-Cnp1 (and Sid4-RFP as SPB marker). Shown are percentages based on analysis of *n* cells (*csi1∆* data is the same as in Fig 2C). **(D)** Model for the impact of SUMOylation on Lem2 function in centromere clustering and silencing.

Finally we questioned whether the function of Lem2 SUMOylation in rescue of centromere clustering in *csi1Δ* cells relates solely to Lem2 localisation, or whether SUMOylation also enhances Lem2 function in centromere clustering independent of localisation. It has been shown previously that Bqt4 functions to mobilise Lem2 away from the SPB and around the nuclear envelope, and that while Lem2 localisation at the SPB is destabilised in *csi1Δ* cells, it is rescued in *bqt4∆ csi1∆* double mutant cells (Ebrahimi et al., 2018). We therefore asked whether the enhanced localisation of Lem2 to the SPB in *bqt4∆ csi1∆* cells would be sufficient to rescue centromere clustering defects, as in *nup132∆ csi1∆* cells. However, this was not the case: while deletion of *bqt4*^*+*^ alone had little effect on centromere clustering, clustering defects in *bqt4∆ csi1∆* double mutant cells were equivalent to those in *csi1∆* single mutant cells (Fig 4C). This suggests that SUMOylation of Lem2 plays an important role in its function in clustering, beyond enhancing its localisation at the SPB.

## Discussion

The dissociation of the deSUMOylase Ulp1 from the nuclear envelope in the absence of Nup132 has been found to cause seemingly paradoxical effects: on the one hand, there are reduced levels of E3 SUMO ligase Pli1 (Nie and Boddy, 2015) and thus reduced global SUMOylation, yet on the other there are toxic effects of persevering polySUMOylated proteins that accumulate at NPCs (Kramarz et al., 2021). Contrary to previous assumptions, here we show that centromere-related defects in *nup132Δ* cells are not explained by destabilisation of Pli1. Rather, we find that suppressing global SUMOylation, or Lem2 SUMOylation specifically, is sufficient to suppress centromeric defects, strongly supporting the model that enhanced SUMOylation at the nuclear periphery is the primary driver of centromeric phenotypes in *nup132∆* cells. Unexpectedly, we also find that while hyper-SUMOylation at the nuclear periphery is detrimental to centromere silencing, it can enhance centromere clustering: clustering defects in *csi1∆* cells are partially rescued by deletion of *nup132*^*+*^, while depleting SUMO suppresses this rescue. We show that Lem2 is again a key SUMO substrate in this context, since specifically suppressing Lem2 SUMOylation is also sufficient to partially supress the rescue. Our results reveal opposing effects of hyper-SUMOylation in general, and Lem2 SUMOylation in particular, on different aspects of centromere function, and suggest a key role for SUMOylation in regulating the diverse activities of Lem2.

### Lem2 SUMOylation may serve as a regulatory switch

Lem2 has been shown to play multiple roles at the nuclear periphery including organisation and silencing of heterochromatin domains, maintenance of nuclear membrane integrity and regulation of nuclear-exosome-mediated RNA degradation (Hiraoka et al., 2011; Gonzalez et al., 2012; Banday et al., 2016; Barrales et al., 2016; Tange et al., 2016; Ebrahimi et al., 2018; Hirano et al., 2018; Pieper et al., 2020; Martin Caballero et al., 2022). Interactions with many different partner proteins are thought to enable Lem2 to localise to distinct subcellular domains and contribute to different pathways. Interestingly, evidence indicates that several of these interactions occur through the same (C-terminal MSC) domain of Lem2 (Gu et al., 2017; Pieper et al., 2020; Martin Caballero et al., 2022), yet how competing interactions are regulated to coordinate distinct functions of Lem2 remains largely unknown. The data presented here suggests that hyper-SUMOylation both impairs Lem2 function in centromere silencing, and simultaneously enhances its function in centromere clustering. As the most parsimonious explanation for these observations, we favour a model in which SUMOylation regulates the competition between Lem2-associating factors, specifically with Lem2 hyper-SUMOylation in *nup132Δ* cells enhancing interactions with centromere clustering components at the cost of those that mediate silencing functions (Fig 4D). Evidence of competition between alternative functions of Lem2 has been observed before: in the related fission yeast *S. japonicus*, the ESCRT-III/Vps4 machinery has been shown to remodel Lem2 heterochromatin attachments, and ESCRT-III/Vps4-mediated release from heterochromatin is required to free up Lem2 to perform its function in nuclear envelope sealing at the end of mitosis (Pieper et al., 2020). We propose that spatially-controlled deSUMOylation of Lem2 represents another layer of regulation of Lem2 interactions, such that Lem2 SUMOylation acts as a regulatory switch between pathways.

Our model proposes a role for SUMOylation in regulating Lem2 function, yet mutation of the seven known SUMOylation sites within Lem2 (Lem2^K7R^) in an otherwise wild-type background does not result in any of the centromere-related phenotypes normally associated with impaired Lem2 function. However, our genetic analyses suggest that this may reflect redundancy with other pathways, since: (1) expressing Lem2^K7R^ in *nup132∆* cells results in increased TBZ sensitivity; and (2) expressing Lem2^K7R^ in *nup132∆ csi1∆* cells causes more severe centromere clustering defects. That the role of Lem2 SUMOylation might be hidden by parallel pathways is consistent with previous findings that show multiple elements of Lem2 function are masked by redundant mechanisms (Barrales et al., 2016).

### Impact of Lem2 SUMOylation on centromeric silencing

We show here that hyper-SUMOylation contributes to centromere silencing defects in *nup132Δ* cells, and that this effect is dependent on SUMOylation of Lem2. Mutation of Lem2 SUMOylation sites rescues the silencing defects without any obvious effect on Lem2 localisation or stability, suggesting that SUMOylation likely alters Lem2 protein-protein interactions. The mechanism by which Lem2 promotes heterochromatic silencing remains opaque; although there is evidence that Lem2 influences the balance of recruitment of opposing chromatin factors including the HDAC repressor complex SHREC and anti-silencing Epe1, no physical association of Lem2 with these factors has been detected (Banday et al., 2016; Barrales et al., 2016). Interestingly, very recent evidence indicates that Lem2 can also influence post-transcriptional silencing by interacting with RNA surveillance factor Red1 to regulate RNA degradation by the nuclear exosome (Martin Caballero et al., 2022). An intriguing possibility is that this post-transcriptional mechanism might contribute to Lem2-mediated silencing of the centromeric *imr:ura4+* reporter; in future it will be interesting to explore whether Lem2 interaction with Red1 is affected by SUMOylation.

An unexpected finding was that suppression of Lem2 SUMOylation can rescue centromere silencing defects in *nup132Δ* cells only when the N-terminus of Lem2 remains intact. This was surprising since the Lem2 N-terminal domain was previously shown to be dispensable for centromeric silencing (Banday et al., 2016; Barrales et al., 2016). A possible explanation is that in *nup132Δ* cells there is hyper-SUMOylation, and therefore potentially misregulation, of one or more other proteins whose function can normally compensate for absence of the Lem2 N-terminal domain. In particular, The N-terminus of Lem2 has been shown to be important for Lem2 binding to centromeric chromatin, yet expression of the Lem2 C-terminus alone is sufficient to maintain centromeric silencing in an otherwise wild-type background (Barrales et al., 2016; Tange et al., 2016). We speculate that in this scenario, interactions with other centromere-localised proteins may be sufficient to localise Lem2 within sufficient proximity to centromeric chromatin to perform its silencing function. However, in *nup132Δ* cells, hyper-SUMOylation of one or more of these proteins might impair Lem2 interactions, such that there is increased dependency on N-terminus-mediated chromatin binding of Lem2 to allow the C-terminus to perform its function in silencing.

### Role of SUMOylation in centromeric clustering

Csi1 and Lem2 have been shown to function in parallel pathways for centromere clustering (Barrales et al., 2016), but dissecting the relative contributions of these pathways is complicated by the fact that Csi1 also plays a role in stabilising Lem2 localisation at the SPB (Ebrahimi et al., 2018). Unexpectedly, here we have found that *nup132*^*+*^ deletion causes a SUMO-dependent rescue in centromere clustering defects in *csi1Δ* cells, and that this is at least partially mediated through SUMOylation of Lem2, since rescue is suppressed upon expression of Lem2^K7R^. Interestingly, the rescue is associated with SUMOylation-dependent enhancement of Lem2 localisation at the SPB. However, deletion of Bqt4 has also been shown to increase Lem2 localisation to the SPB, even in the absence of Csi1 (Ebrahimi et al., 2018), yet we find that *bqt4Δ csi1Δ* cells show no alleviation in centromere clustering defects. Together these observations suggest that deletion of *nup132*^*+*^ causes increased SUMOylation of Lem2 that enhances its localisation at SPB in *csi1Δ* cells, however this localisation is not sufficient to rescue centromere clustering defects, and hyper-SUMOylation plays an additional role in enhancing centromere clustering adjacent to the SPB.

How might enhanced SUMOylation influence centromere clustering pathways? Hyper-SUMOylated proteins can be targeted by STUbL complexes resulting in ubiquitin-dependent extraction and/or proteasomal degradation, as seen for Pli1 in *nup132Δ* cells (Uzunova et al., 2007; Nie and Boddy, 2015; Nie and Boddy, 2016). However, analyses in *slx8Δ* and *ufd1∆Ct*^*213-342*^ backgrounds revealed no evidence that enhancement of centromere clustering in *nup132Δ* cells involves targeting of SUMOylated substrates *via* STUbL/ufd1 pathways. We therefore speculate that a SUMO “molecular glue” mechanism may be enhancing centromere clustering, similar to what has been described for Promyelocytic leukemia nuclear bodies (Shen et al., 2006; Corpet et al., 2020). PML-NBs are membrane-free compartments that regulate a number of processes including transcriptional control and DNA repair, and importantly both SUMOylation of PML and internal SUMO interaction motifs (SIMs) are key in NB formation through non-covalent SUMO-SIM interactions. We hypothesise that SUMOylated Lem2 may interact with as yet undiscovered SIM domains within Lem2-associated proteins, enhancing both localisation of Lem2 to the SPB and recruitment of centromere clustering factors. Of note, whilst PML-NB formation is mediated by SUMOylation, the process is also driven by liquid-liquid phase separation in intrinsically disordered proteins (Banani et al., 2017; Shin and Brangwynne, 2017; Zhang et al., 2020). Strikingly, LEM2 has been observed to form liquid-liquid phase droplets in human cells (Gu et al., 2017); an unexplored question is whether *S. pombe* Lem2 also has the ability to phase-separate, and whether SUMOylation of Lem2 could mediate such processes, similar to regulation of PML-NBs.

### Other possible SUMO substrates of relevance to centromere function

We observe that expression of Lem2^K7R^ is sufficient to partially but not fully suppress the *nup132Δ*-mediated rescue of centromere clustering in *csi1Δ* cells. One possible explanation for this is that Lem2^K7R^ can still be SUMOylated *via* remaining lysines, albeit presumably to a lesser degree. This could reflect presence of other physiological SUMOylation sites not identified in previous analyses, or targeting of other lysines for SUMO-modification in the absence of usual physiological sites. In an attempt to more comprehensively suppress Lem2 SUMOylation, we generated strains in which Lem2 was fused to the catalytic domain of SUMO protease Ulp1, a strategy successfully employed previously to inhibit SUMOylation of Top2 (Wei et al., 2017). However, we found that fusion of either the active or inactive catalytic domain of Ulp1 impaired Lem2 function, precluding further analysis. Therefore the question remains as to whether complete suppression of Lem2 SUMOylation would fully prevent the rescue of centromere clustering.

Another, non-mutually exclusive, explanation for why the effects of *nup132*^*+*^ deletion are not fully suppressed by expression of Lem2^K7R^ is that SUMOylation of one or more additional proteins also contributes to the effects. Factors related to centromeres and the nuclear periphery are highly enriched amongst SUMOylated proteins (Kohler et al., 2015); Lem2 may therefore be one of several proteins whose increased SUMOylation can contribute to enhanced centromere clustering in circumstances where this process is impaired. Perhaps related to this, it has been shown previously that deletion of *nup132*^*+*^ disrupts the normal dynamic disassembly and reassembly of the outer kinetochore during meiotic prophase in *S. pombe* (Yang et al., 2015); it is tempting to speculate that this may also be caused by constitutive hyper-SUMOylation of component proteins due to the loss of localised Ulp1-mediated deSUMOylation.

We note that while suppressing Lem2 SUMOylation (Lem2^K7R^) rescues centromere silencing defects in *nup132Δ* cells, it conversely exacerbates TBZ sensitivity in this background. This is despite the fact that reducing global polySUMOylation through overexpression of Ulp1 or Pmt3^KallR^ alleviates TBZ sensitivity. This suggests that while SUMOylation of Lem2 plays at least a partial role in both reducing pericentromeric silencing and simultaneously enhancing centromere clustering, hyper-SUMOylation of further unidentified protein(s) is likely responsible for TBZ sensitivity. Identifying these proteins may be challenging, for example if multiple factors contribute redundantly. However, TBZ sensitivity can be caused by reduced accumulation of cohesins at centromeres, and interestingly, studies in human cell culture have revealed that hyper-SUMOylation of RAD21 and other cohesin subunits reduces their chromatin association (Wagner et al., 2019). *S. pombe* Rad21 has also been found to be SUMOylated (Kohler et al., 2015), hence this could be a promising candidate for a protein whose hyper-SUMOylation could contribute to TBZ sensitivity in *nup132Δ* cells. Levels of human RAD21 SUMOylation are regulated through the specific localisation of the deSUMOylating enzyme SENP6 at centromeric and telomeric domains (Wagner et al., 2019). Given the localisation of centromeres to the nuclear envelope in fission yeast, an attractive model is that only centromere-localised Rad21 that is within proximity of nuclear envelope-tethered Ulp1 is deSUMOylated, allowing centromere-specific deSUMOylation and enhanced chromatin binding at specific loci.

### Concluding Remarks

Through analysis of *nup132Δ* cells, we have found that hyper-SUMOylation at the nuclear periphery both impairs centromeric silencing, and enhances clustering of centromeres at the SPB. Both physical clustering (Muller et al., 2019), and enrichment for SUMOylated proteins (Abrieu and Liakopoulos, 2019), appear to be common features of centromeres across diverse species, while changes in centromere clustering have been linked to human carcinoma progression (Verrelle et al., 2021), yet mechanisms of clustering remain poorly defined. It will be interesting to see whether SUMO may play a conserved role in promoting centromere clustering, possibly acting as a “molecular glue” to facilitate protein-protein interactions. In addition, we show that Lem2 is a key SUMO substrate in the context of both centromere silencing and clustering, and present a novel model whereby SUMOylation may play an important role in modulating the balance of Lem2 interactions with partner proteins to coordinate its diverse functions. Mammalian Lem2 is critical for embryonic development (Tapia et al., 2015), and mutations in LEM2 have been linked to several human diseases including juvenile cataracts, arrhythmic cardiomyopathy, and a novel nuclear envelopathy with progeria-like symptoms (Boone et al., 2016; Abdelfatah et al., 2019; Marbach et al., 2019), highlighting the potential clinical importance of understanding the regulation of Lem2 mediated pathways. In *S. pombe*, it has previously been suggested that post-translational modification could account for observed differences in Lem2 function in different nutritional conditions (Martin Caballero et al., 2022); it is an intriguing possibility that changes in Lem2 SUMOylation status may mediate the response to environmental cues.

## Materials and Methods

### Yeast strains and plasmids

Detailed lists of strains and plasmids are provided in Tables S2 and S3, respectively. For deletion and epitope tagging (with Flag or GFP), a PCR-based method was used to amplify resistance cassettes flanked by 80bp target site homology for integration at endogenous loci by homologous recombination (Bahler et al., 1998). For Pli1^K3R^ and Lem2^K7R^ strains, long homology-containing fragments incorporating the relevant lysine to arginine mutations were generated by fusion PCR, and integrated at endogenous loci by transformation into *pli1Δ::ura4*^*+*^ or *lem2Δ::ura4*^*+*^ strains, respectively, followed by selection on 5-FOA. For over-expression of a mature Pmt3 construct that does not require processing by Ulp1, a pREP41-myc-his-pmt3^+^ plasmid (Jongjitwimol et al., 2014) was modified by site directed mutagenesis to insert a stop codon immediately after residue G111 (resulting in terminal di-glycine). The Pmt3^KallR^ over-expression plasmid was subsequently generated from the pREP41-myc-his-mature-pmt3^+^ plasmid by further rounds of site-directed mutagenesis. To generate the Ulp1 over-expression plasmid, genomic *ulp1*^*+*^ was amplified by PCR and integrated into pREP41-myc-his plasmid using Gibson assembly methods. Genomic modifications and plasmids were verified by sequencing.

Cells were grown in YES medium, with the exception of strains expressing pREP41 plasmids which were maintained in PMG -Leu. For spotting assays, 10-fold serial dilutions were plated onto non-selective media, or media supplemented with 1 g/L 5-FOA, or 20 µg/mL TBZ.

### Global analysis of ubiquitination sites

Enrichment for ubiquitinated peptides using PTMScan® beads broadly followed the detailed protocol published previously (Udeshi et al., 2013), with the following exceptions to apply the method in *S. pombe* (a broad outline is shown in Fig S1A). Approximately 3 × 10^9^ wild-type *S. pombe* cells were grown to mid-log phase in YES liquid, washed twice with PBS and harvested. Cells were lysed by bead-beating in a denaturing lysis buffer (8 M Urea, 50 mM Tris-HCl (pH 8.0), 150 mM NaCl, 1 mM EDTA, 6 mM MgCl_2_, 1 mM PMSF, 4 mM NEM, 50 µM PR-619, 20 µM MG-132, 4 mM 1, 10-phenanthroline, 1× complete EDTA-free protease inhibitor cocktail (Roche) and 1 mM pefabloc) using acid washed beads and a VXR basic Vibrax® at 4 °C. The lysate was clarified by centrifugation at 17000 rcf for 30 mins at 4 °C, and a Bradford assay was performed to assess protein concentration. Reduction, alkylation, LysC and trypsin digestion steps were performed as described previously (Udeshi et al., 2013). Peptide reverse phase offline fractionation was performed on a Dionex Ultimate 5000 HPLC (Thermo Scientific) as described previously (Udeshi et al., 2013) with minimal variations. In brief, peptides were loaded on a Zorbax 300-Extend-C-18 (5 µm, 4.6 × 250 mm) column (Agilent) at a constant flow rate of 1 mL/min. Peptides were separated according to the following gradient.

| Retention (min) | Flow<br>(mL/min) | %B |
| --- | --- | --- |
| 0.00 | 1.000 | 0.00 |
| 5.00 | 1.000 | 3.00 |
| 7.00 | 1.000 | 5.00 |
| 35.00 | 1.000 | 15.00 |
| 45.00 | 1.000 | 20.00 |
| 53.00 | 1.000 | 35.00 |
| 60.00 | 1.000 | 50.00 |
| 62.00 | 1.000 | 70.00 |
| 65.00 | 1.000 | 70.00 |
| 70.00 | 1.000 | 0.00 |
| 82.00 | 1.000 | 0.00 |

Fractions were collected at 1 min time slices until the 64^th^ minute, and vacuum centrifuged to dryness. Dried peptides were resuspended in 187.5 µL IAP buffer (50 mM MOPS-pH 7.2, 10 mM sodium phosphate, 50 mM NaCl), and fractions were pooled in a serpentine, non-contiguous manner such that every 8^th^ fraction was combined to generate 8 final fractions containing 1.5 mL of resuspended peptides. DiGly remnant enrichment using PTMScan® beads was performed as described previously (Udeshi et al., 2013).

### LC-MS/MS analysis

LC-MS/MS analyses were performed on Q Exactive Plus and Q Exactive mass spectrometers (both Thermo Scientific). For the Q Exactive Plus analysis, liquid chromatography for the LC-MS/MS runs was performed on an EASY-nLC 1000 liquid chromatography system (Thermo Scientific) coupled to spectrometers *via* modified NanoFlex sources (Thermo Scientific).

Peptides were loaded onto 250 mm × 75 µm PicoFrit (C18, 2 µm medium) analytical columns (New Objective) at a maximum pressure of 800 bar. Solutions A and B for ultra-performance LC were 0.1% FA in water and ACN, respectively. Peptides were eluted into the mass spectrometer at a flow rate of 200 nL/min using a gradient that incorporated a linear phase from 6% B to 30% B in 80 min, followed by a steeper phase and wash. The gradient run time was ∼120 min. The Q Exactive Plus mass spectrometer was operated in the data-dependent mode acquiring HCD MS/MS scans in the 300-1,800 m/z scan range with a resolution of 17,500 after each MS1 scan (Resolution= 70,000) on the 12 most abundant ions using an MS1 ion target of 3 × 10^6^ ions and an MS2 target of 5 × 10^5^ ions. The maximum ion time utilized for MS/MS scans was 120 ms; the HCD-normalized collision energy was set to 25 and the dynamic exclusion time was set to 20 s. The peptide match and isotope exclusion functions were enabled. The Q Exactive mass spectrometer (Thermo Fisher Scientific) was coupled on-line to a 50 cm Easy-Spray column (Thermo Fisher Scientific), which was assembled on an Easy-Spray source and operated constantly at 50 °C. Mobile phase A consisted of 0.1% formic acid, while mobile phase B consisted of 80% acetonitrile and 0.1% formic acid. Peptides were loaded onto the column at a flow rate of 0.3 μL/min and eluted at a flow rate of 0.25 μL/min according to the following gradient: 2 to 40% buffer B in 120 min, then to 95% in 11 min (total run time of 160min). Survey scans were performed at 70,000 resolution (scan range 350-1400 m/z) with an ion target of 1 × 10^6^ and injection time of 20ms. MS2 was performed with an ion target of 5 × 10^4^, injection time of 60 ms and HCD fragmentation with normalized collision energy of 27. The isolation window in the quadrupole was set at 2.0 Thomson. Only ions with charge between 2 and 7 were selected for MS2.

## Data analysis

The MaxQuant software platform (Cox and Mann, 2008) version 1.6.1.0 was used to process the raw files and search was conducted against the *Schizosaccharomyces pombe* (July, 2016) protein database, using the Andromeda search engine (Cox et al., 2011). For the first search, peptide tolerance was set to 20 ppm while for the main search peptide tolerance was set to 4.5 pm. Isotope mass tolerance was 2 ppm and maximum charge to 7. Digestion mode was set to specific with trypsin allowing maximum of two missed cleavages. Carbamidomethylation of cysteine was set as fixed modification. Oxidation of methionine, and the diGly residue on lysine were set as variable modifications. Peptide and protein identifications were filtered to 1% FDR.

### RT-qPCR

Total RNA was extracted from 1 × 10^7^ mid-log phase cells using the Masterpure Yeast RNA Purification Kit (Epicentre), according to the manufacturer’s instructions. 1 µg of extracted RNA was treated with TURBO DNase (Ambion) for 1 hour at 37 °C, and reverse transcription was performed using random hexamers (Roche) and Superscript III reverse transcriptase (Invitrogen). Lightcycler 480 SYBR Green (Roche) and primers (_q_act1_F: GTTTCGCTGGAGATGATG; _q_act1_R: ATACCACGCTTGCTTTGAG; _q_ura4_F: CGTGGTCTCTTGCTTTGG; _q_ura4_R: GTAGTCGCTTTGAAGGTTAGG) were used for qPCR quantification of *imr:ura4*^*+*^ transcript levels for relative to *act1*^*+*^. Data presented represent three biological replicates and error bars represent one standard deviation. P-values were calculated using Student’s *t* test.

### Immunoprecipitation

Immunoaffinity purifications were performed essentially as previously described (Oeffinger et al., 2007). Cultures were grown to mid-log phase in YES and 3 × 10^8^ cells were harvested in either (50 mM Tris-HCl (pH 8.0), 150 mM NaCl, 1 mM EDTA, 1 mM PMSF, 1× complete EDTA-free protease inhibitor cocktail (Roche)) for Pli1-Flag IP, or (50 mM Hepes-pH 7.5, 150 mM NaCl, 1% Triton X100, 1 mM EDTA, 2 mM PMSF, 1× complete EDTA-free protease inhibitor cocktail (Roche)) for Lem2-GFP IP. Cells were lysed by bead beading, and supernatant was clarified by 2 × 10 mins centrifugation at 17000 rcf at 4 °C. Extracts were incubated with pre-equilibrated protein G agarose and anti-Flag M2 (Merck), or protein A agarose and anti-GFP (A-11122 Thermo), for 1 hour at 4 °C. Beads were washed three times with lysis buffer, and proteins eluted in gel loading buffer (150 mM Tris, 8M urea, 2.5% (w/v) SDS, 20% (v/v) glycerol, 10% (w/v) 2-mercaptoethanol, 3% (w/v) DTT, 0.1% (w/v) bromophenol blue, pH 6.8(HCl)) and analysed by immunoblotting using anti-Flag M2 (Merck, 1:1000 dilution), anti-GFP (A-11122 Thermo, 1:1000 dilution) and anti-Tat1 (mouse monoclonal anti-tubulin from Keith Gull, 1:200 dilution).

### Live cell imaging

Cells expressing Lem2-GFP, GFP-Cnp1 and Sid4-RFP were grown to mid-log phase in YES (except for those strains expressing plasmids which were grown in PMG -Leu) and embedded in low-melting point agarose. Imaging was performed at 25 °C using a Nikon Ti2 inverted microscope, equipped with a 100x 1.49 NA Apo TIRF objective and a Teledyne Photometrics Prime 95B camera. Images were acquired with NIS-elements (version 5.1), with z-stacks taken at 0.25 µm intervals. Maximum intensity Z-projections were made in ImageJ. For centromere clustering analyses, chi-squared (χ^2^) tests were performed to calculate p-values for differences in the proportions of cells displaying centromeres ‘clustered’ *vs* ‘unclustered’ (1 GFP-Cnp1 focus *vs* 2/3 GFP-Cnp1 foci). Similarly, for analysis of Lem2 localisation, χ^2^ tests were used to calculate p-values for differences in the proportion of cells in which Lem2 was ‘present’ versus ‘absent’ at the SPB (denoted by Sid4-RFP).

## Supplementary material

Table S1 lists the identified ubiquitination sites. Table S2 lists the yeast strains used in the study. Table S3 lists the plasmids used in the study.

## Acknowledgements

We thank Yasushi Hiraoka, Robin Allshire, Genevieve Thon, Matthew Whitby and Julia Cooper for strains, Felicity Watts for the Pmt3 expression plasmid, Keith Gull for the anti-Tat1 antibody and Ivan Matic for assistance with proteomic analyses. This work was supported by a Wellcome Trust Investigator Award (202771/Z/16/Z), UK Medical Research Council Career Development Award (G1000505) and Leverhulme Trust project grant (RPG-2014-050) to EHB. Imaging was performed in Centre Optical Instrumentation Laboratory (COIL), which is supported by a Core Grant (203149) to the Wellcome Centre for Cell Biology at the University of Edinburgh.

The authors declare no competing financial interests.

## Author contributions

Conceptualisation: J. Strachan, O. Leidecker and E.H. Bayne. Methodology: J. Strachan. O. Leidecker, C Spanos, E. Chapman and E.H. Bayne. Investigation: J. Strachan, O. Leidecker, C Spanos, C. Le Coz, E. Chapman, A. Arsenijevic, H. Zhang, N. Zhao. Data Curation: J. Strachan. O. Leidecker, C Spanos. Formal Analysis: J. Strachan. O. Leidecker, C Spanos, C. Le Coz. Writing – original draft: J. Strachan and E.H. Bayne. Writing – review and editing: all authors. Visualisation: J. Strachan and E.H. Bayne. Supervision: E.H. Bayne. Funding acquisition: E.H. Bayne.

---

**Figure S1.**
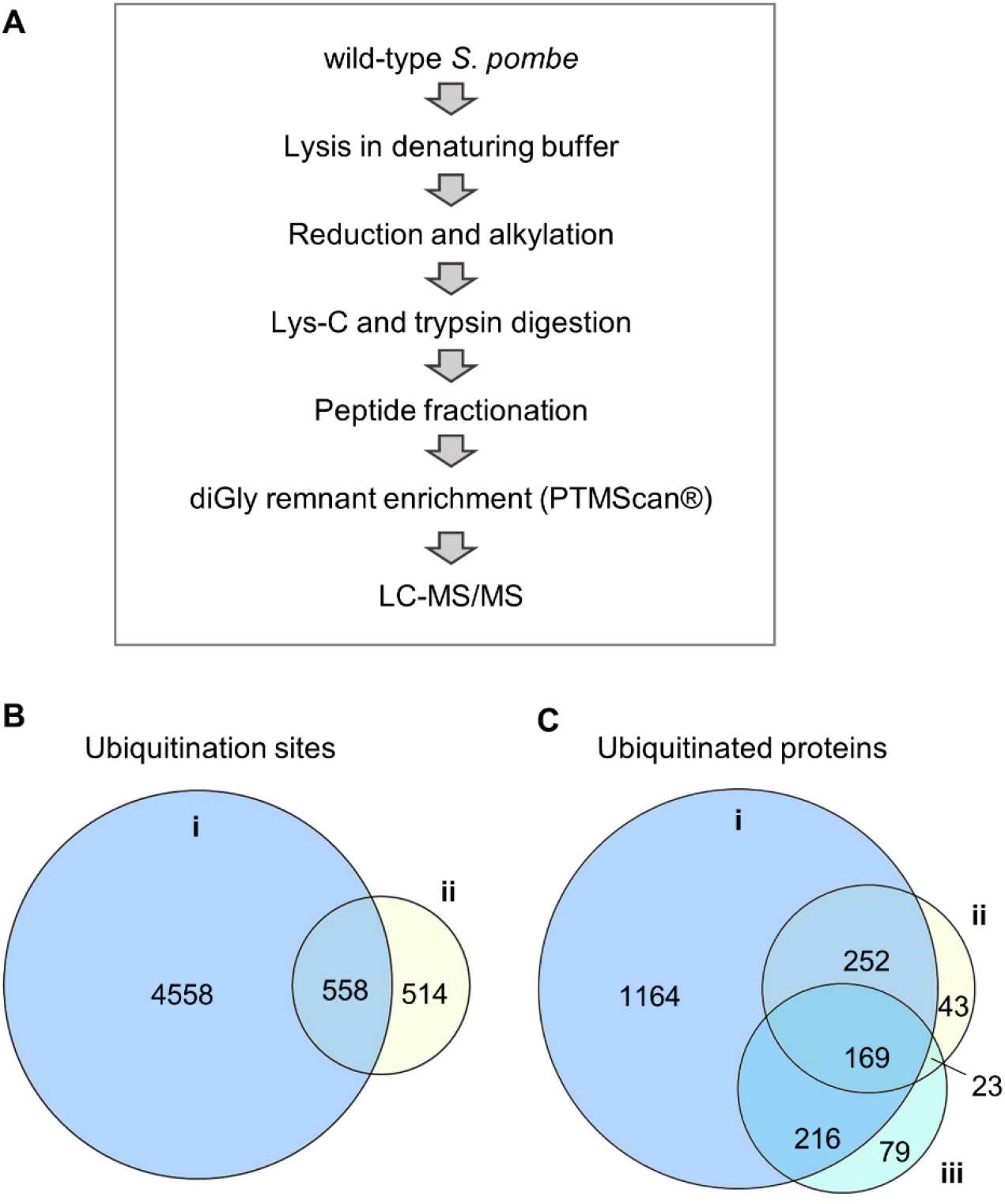
Global analysis of ubiquitination sites in fission yeast. **(A)** Outline of the approach employed to identify ubiquitination sites under physiological conditions. **(B)** Comparison of the number of ubiquitination sites identified in this study (i) versus the previous study (Beckley et al., 2015) (ii). 558 sites identified previously were found in this analysis, equating to ∼55% of the total ubiquitination sites previously identified. **(C)** Comparison of the number of ubiquitinated proteins identified in this study (i) versus the previous study (Beckley et al., 2015). The latter are divided into those identified directly by identification of diGly peptides (ii); and those identified indirectly based on altered abundance in the absence of deubiquitinating enzymes (iii). 169 proteins were identified in all three analyses.

**Figure S2.**
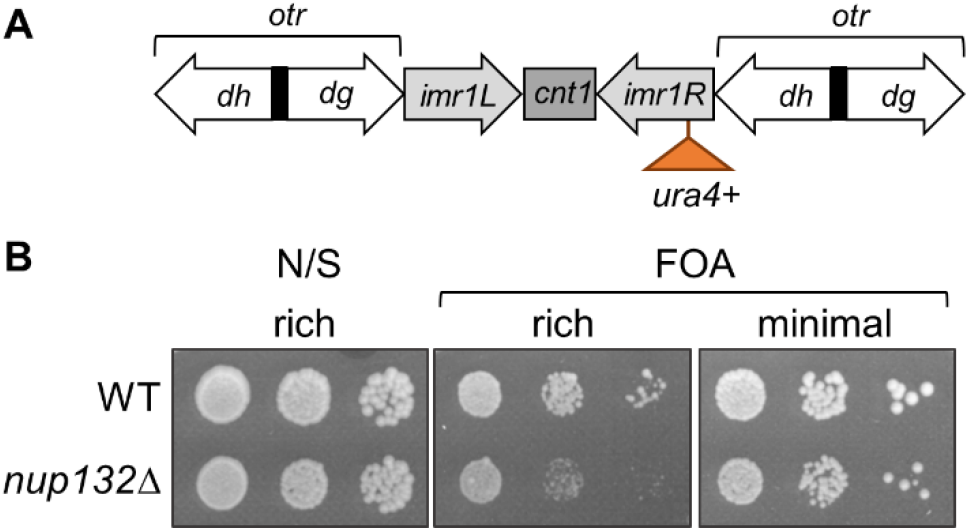
Deletion of *nup132*^*+*^ results in defects in centromeric silencing in rich media but not minimal media. **(A)** Schematic representation of the *imr1:ura4*^*+*^ reporter, indicating the position of the *ura4*^*+*^ insertion in centromere one relative to centromeric outer repeats (*otr*; *dg* and *dh*), innermost repeats (*imr*) and central core (*cnt*). **(B)** Assay for silencing of the *imr1:ura4*^*+*^ reporter in rich (YES) or minimal (PMG) media: loss of silencing results in increased expression of *ura4*^*+*^ and therefore decreased growth in the presence of the counter-selective drug 5-FOA. Growth in the absence of 5-FOA (non-selective, N/S) is shown as a control.

**Figure S3.**
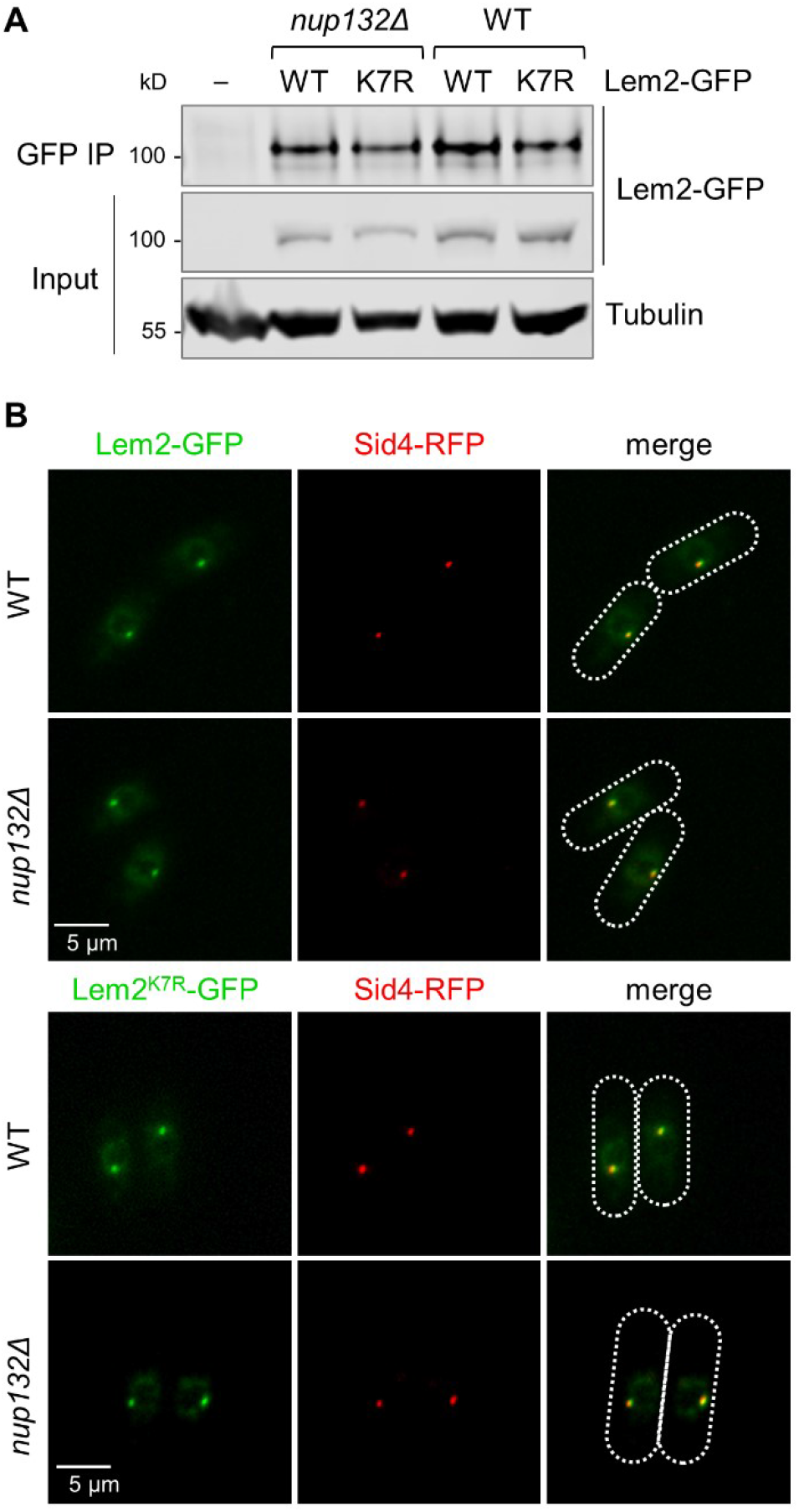
Lem2 stability and localisation are unaffected by mutation of Lem2 localisation sites or deletion of *nup132*^*+*^. **(A)** Western blot analysis of Lem2-GFP, or Lem2^K7R^-GFP, immunoprecipitated from wild-type or *nup132∆* cells. Tubulin (anti-Tat1) serves as a loading control. **(B)** Representative images from two-colour live cell imaging of Lem2-GFP, or Lem2^K7R^-GFP, and Sid4-RFP (SPB marker) in wild-type and *nup132∆* cells. Dotted lines indicate cell boundaries.

**Figure S4.**
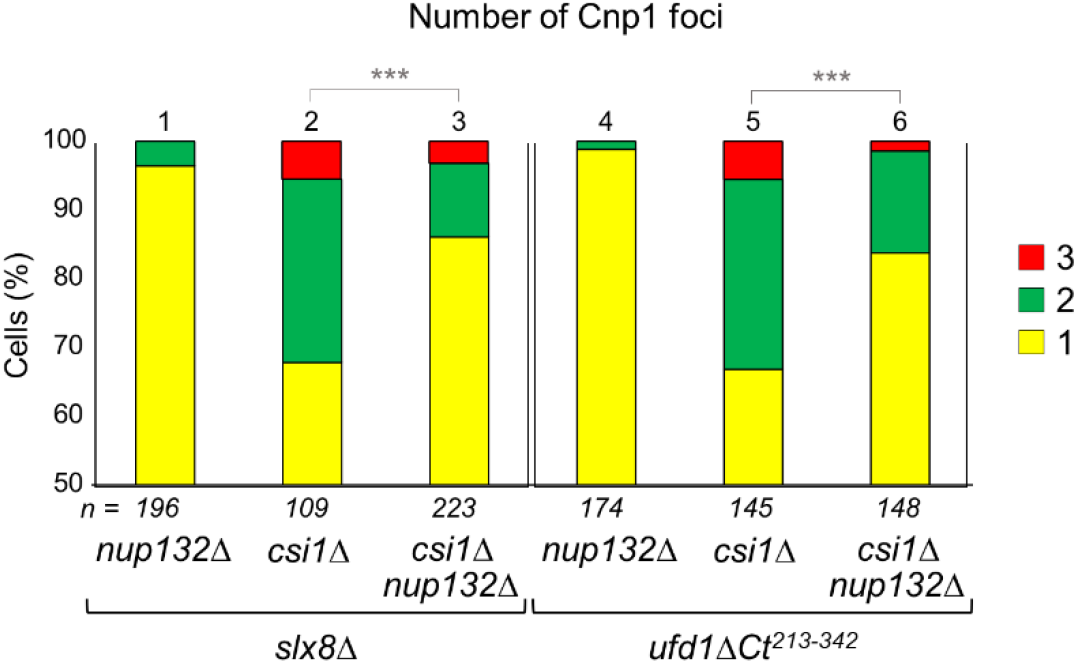
SUMO-mediated enhancement of centromere clustering is not dependent on Slx8 or Ufd1. Quantification of cells displaying one, two or three Cnp1 foci, based on live-cell imaging of GFP-Cnp1 (and Sid4-RFP as SPB marker). Shown are percentages based on analysis of *n* cells. Asterisks (***) denote *p* ≤ 0.001 from χ2 test analysis.

**Supplementary Table S2.** Strains used in this study.

| Strain | Genotype | Figure |
| --- | --- | --- |
| 3381 | *h+ ade6-210 arg3-D4 his3-D1 leu1-32 ura4-D18 otr1R(dg-glu)Sph1:ade6<sup>+</sup>* | 1b |
| 3998 | *h- pli1<sup>+</sup>-flag:NatR ade6-210 arg3-D4 his3-D1 leu1-32 ura4-D18 otr1R(dg-glu)Sph1:ade6<sup>+</sup>* | 1b |
| 3999 | *h- pli1<sup>K3R</sup>-flag:NatR ade6-210 arg3-D4 his3-D1 leu1-32 ura4-D18 otr1R(dg-glu)Sph1:ade6<sup>+</sup>* | 1b |
| 4002 | *h+ nup132Δ::ura4<sup>+</sup> pli1<sup>+</sup>-flag:NatR ade6-210 arg3-D4 his3-D1 leu1-32 ura4-D18 otr1R(dg-glu)Sph1:ade6<sup>+</sup>* | 1b |
| 4003 | *h- nup132Δ::ura4<sup>+</sup> pli1<sup>K3R</sup>-flag:NatR ade6-210 arg3-D4 his3-D1 leu1-32 ura4-D18/DSE otr1R(dg-glu)Sph1:ade6<sup>+</sup>* | 1b |
| 4371 | *clr4Δ::leu2<sup>+</sup> ade6-210 leu1-32 ura4-D18/DSE imr1L(NcoI):ura4<sup>+</sup>* | 1c |
| 737 | *h- leu1-32 ura4DS/E ade6-210 his1-102 imr1L(NcoI):ura4<sup>+</sup>* | 1c, 2d, 2e, 2f, S2, S3a |
| 4252 | *pli1<sup>+</sup>-flag:NatR ade6-210 arg3-D4 his3-D1 leu1-32 ura4-D18 imr1L(NcoI):ura4<sup>+</sup>* | 1c |
| 4259 | *pli1<sup>K3R</sup>-flag:NatR ade6-210 arg3-D4 his3-D1 leu1-32 ura4-D18/DSE imr1L(NcoI):ura4<sup>+</sup>* | 1c |
| 4365 | *h+ nup132Δ::HygR ade6-210 arg3-D4 his3-D1 leu1-32 ura4-D18/DSE imr1L(NcoI):ura4<sup>+</sup>* | 1c, 2d, 2e, 2f, S2 |
| 4368 | *pli1Δ::NatR ade6-210 arg3-D4 his3-D1 leu1-32 ura4-D18/DSE imr1L(NcoI):ura4<sup>+</sup>* | 1c |
| 4330 | *nup132Δ::HygR pli1<sup>+</sup>-flag:NatR ade6-210 arg3-D4 his3-D1 leu1-32 ura4-D18 imr1L(NcoI):ura4<sup>+</sup>* | 1c |
| 4370 | *nup132Δ::HygR pli1<sup>K3R</sup>-flag:NatR ade6-210 arg3-D4 his3-D1 leu1-32 ura4-D18/DSE imr1L(NcoI):ura4<sup>+</sup>* | 1c |
| 5192 | *pREP41 ade6-210 leu1-32 ura4-D18/DSE imr1L(NcoI):ura4<sup>+</sup>* | 2a |
| 5193 | *pREP41 ade6-210 leu1-32 ura4-D18/DSE imr1L(NcoI):ura4<sup>+</sup>* | 2a |
| 5158 | *pREP41-myc-his-ulp1<sup>+</sup> ade6-210 leu1-32 ura4-D18/DSE imr1L(NcoI):ura4<sup>+</sup>* | 2a |
| 5159 | *pREP41-myc-his-ulp1<sup>+</sup> ade6-210 leu1-32 ura4-D18/DSE imr1L(NcoI):ura4<sup>+</sup>* | 2a |
| 4984 | *pREP41 nup132Δ::HygR ade6-210 arg3-D4 his3-D1 leu1-32 ura4-D18/DSE imr1L(NcoI):ura4<sup>+</sup>* | 2a |
| 4985 | *pREP41 nup132Δ::HygR ade6-210 arg3-D4 his3-D1 leu1-32 ura4-D18/DSE imr1L(NcoI):ura4<sup>+</sup>* | 2a |
| 4987 | *pREP41-myc-his-ulp1<sup>+</sup> nup132Δ::HygR ade6-210 arg3-D4 his3-D1 leu1-32 ura4-D18/DSE imr1L(NcoI):ura4<sup>+</sup>* | 2a |
| 4988 | *pREP41-myc-his-ulp1<sup>+</sup> nup132Δ::HygR ade6-210 arg3-D4 his3-D1 leu1-32 ura4-D18/DSE imr1L(NcoI):ura4<sup>+</sup>* | 2a |
| 8894 | *h- pREP41 leu1-32 ura4DS/E ade6-210 his1-102 imr1L(NcoI):ura4<sup>+</sup>* | 2b |
| 8895 | *h- pREP41-myc-his-mature-pmt3<sup>+</sup> leu1-32 ura4DS/E ade6-210 his1-102 imr1L(NcoI):ura4<sup>+</sup>* | 2b |
| 8896 | *h- pREP41-myc-his-mature-pmt3<sup>KallR</sup> leu1-32 ura4DS/E ade6-210 his1-102 imr1L(NcoI):ura4<sup>+</sup>* | 2b |
| 8897 | *h+ pREP41 nup132Δ::HygR ade6-210 arg3-D4 his3-D1 leu1-32 ura4-D18/DSE imr1L(NcoI):ura4<sup>+</sup>* | 2b |
| 8898 | *h+ pREP41-myc-his-mature-pmt3<sup>+</sup> nup132Δ::HygR ade6-210 arg3-D4 his3-D1 leu1-32 ura4-D18/DSE imr1L(NcoI):ura4<sup>+</sup>* | 2b |
| 8899 | *h+ pREP41-myc-his-mature-pmt3<sup>KallR</sup> nup132Δ::HygR ade6-210 arg3-D4 his3-D1 leu1-32 ura4-D18/DSE imr1L(NcoI):ura4<sup>+</sup>* | 2b |
| 5820 | *lem2<sup>K7R</sup> leu1-32 ura4-D18/DSE imr1L(NcoI):ura4<sup>+</sup>* | 2d, 2e |
| 5821 | *lem2<sup>K7R</sup> leu1-32 ura4-D18/DSE imr1L(NcoI):ura4<sup>+</sup>* | 2d |
| 5822 | *nup132Δ::HygR lem2<sup>K7R</sup> leu1-32 ade6<sup>+</sup> ura4-D18/DSE imr1L(NcoI):ura4<sup>+</sup>* | 2d, 2e |
| 5182 | *lem2ΔN nup132Δ::HygR leu1-32 ura4-D18/DSE imr1L(NcoI):ura4<sup>+</sup>* | 2f |
| 5184 | *lem2ΔN leu1-32 ura4-D18/DSE imr1L(NcoI):ura4<sup>+</sup>* | 2f |
| 5513 | *h90 sid4<sup>+</sup>-mRFP:KanR GFP-cnp1<sup>+</sup>:NatR leu1-32 ura4-D18* | 3a, 3b |
| 5517 | *h90 lem2Δ::ura4<sup>+</sup> sid4<sup>+</sup>-mRFP:KanR GFP-cnp1<sup>+</sup>:NatR leu1-32 ura4-D18* | 3a, 3b |
| 5516 | *h- nup132Δ::HygR sid4<sup>+</sup>-mRFP:KanR GFP-cnp1<sup>+</sup>:NatR leu1-32 ura4-D18* | 3a, 3b |
| 6209 | *lem2<sup>K7R</sup> sid4<sup>+</sup>-mRFP:KanR GFP-cnp1<sup>+</sup>:NatR leu1-32 ura4-D18* | 3b |
| 6207 | *lem2<sup>K7R</sup> nup132Δ::HygR sid4<sup>+</sup>-mRFP:KanR GFP-cnp1<sup>+</sup>:NatR leu1-32 ura4-D18* | 3b |
| 6363 | *h+ csi1Δ::HygR GFP-cnp1<sup>+</sup>:NatR sid4<sup>+</sup>-mRFP:KanR leu1-32 ura4-D18* | 3c, 4c |
| 6862 | *csi1Δ::HygR nup132Δ::ura4<sup>+</sup> GFP-cnp1<sup>+</sup>:NatR sid4<sup>+</sup>-mRFP:KanR leu1-32 ura4-D18* | 3c |
| 7036 | *pli1Δ::ura4<sup>+</sup> sid4<sup>+</sup>-mRFP:KanR GFP-cnp1<sup>+</sup>:NatR leu1-32 ura4-D18* | 3c |
| 7015 | *pli1Δ::ura4<sup>+</sup> nup132Δ::HygR sid4<sup>+</sup>-mRFP:KanR GFP-cnp1<sup>+</sup>:NatR leu1-32 ura4-D18* | 3c |
| 7302 | *h- csi1Δ::KanR pli1Δ::ura4<sup>+</sup> sid4<sup>+</sup>-mRFP:KanR GFP-cnp1<sup>+</sup>:NatR leu1-32 ura4-D18* | 3c |
| 7233 | *h- csi1Δ::KanR nup132Δ::HygR pli1Δ::ura4<sup>+</sup> sid4<sup>+</sup>-mRFP:KanR GFP-cnp1<sup>+</sup>:NatR leu1-32 ura4-D18* | 3c |
| 7440 | *h+ csi1Δ::HygR lem2<sup>K7R</sup> GFP-cnp1<sup>+</sup>:NatR sid4<sup>+</sup>-mRFP:KanR leu1-32 ura4-D18/DSE* | 3c |
| 7444 | *h- csi1Δ::HygR nup132Δ::HygR lem2<sup>K7R</sup> GFP-cnp1<sup>+</sup>:NatR sid4<sup>+</sup>-mRFP:KanR leu1-32 ura4-D18/DSE* | 3c |
| 7311 | *h90 pREP41 sid4<sup>+</sup>-mRFP:KanR GFP-cnp1<sup>+</sup>:NatR leu1-32 ura4-D18* | 3d |
| 7315 | *h90 pREP41-myc-his-mature-pmt3<sup>+</sup> sid4<sup>+</sup>-mRFP:KanR GFP-cnp1<sup>+</sup>:NatR leu1-32 ura4-D18* | 3d |
| 7317 | *h90 pREP41-myc-his-mature-pmt3<sup>KallR</sup> sid4<sup>+</sup>-mRFP:KanR GFP-cnp1<sup>+</sup>:NatR leu1-32 ura4-D18* | 3d |
| 7313 | *h90 pREP41-myc-his-ulp1<sup>+</sup> sid4<sup>+</sup>-mRFP:KanR GFP-cnp1<sup>+</sup>:NatR leu1-32 ura4-D18* | 3d |
| 7319 | *h- pREP41 nup132Δ::HygR sid4<sup>+</sup>-mRFP:KanR GFP-cnp1<sup>+</sup>:NatR leu1-32 ura4-D18* | 3d |
| 7323 | *h- pREP41-myc-his-mature-pmt3<sup>+</sup> nup132Δ::HygR sid4<sup>+</sup>-mRFP:KanR GFP-cnp1<sup>+</sup>:NatR leu1-32 ura4-D18* | 3d |
| 7325 | *h- pREP41-myc-his-mature-pmt3<sup>KallR</sup> nup132Δ::HygR sid4<sup>+</sup>-mRFP:KanR GFP-cnp1<sup>+</sup>:NatR leu1-32 ura4-D18* | 3d |
| 7321 | *h- pREP41-myc-his-ulp1<sup>+</sup> nup132Δ::HygR sid4<sup>+</sup>-mRFP:KanR GFP-cnp1<sup>+</sup>:NatR leu1-32 ura4-D18* | 3d |
| 7327 | *h+ pREP41 csi1Δ::HygR GFP-cnp1<sup>+</sup>:NatR sid4<sup>+</sup>-mRFP:KanR leu1-32 ura4-D18* | 3d |
| 7331 | *h+ pREP41-myc-his-mature-pmt3<sup>+</sup> csi1Δ::HygR GFP-cnp1<sup>+</sup>:NatR sid4<sup>+</sup>-mRFP:KanR leu1-32 ura4-D18* | 3d |
| 7333 | *h+ pREP41-myc-his-mature-pmt3<sup>KallR</sup> csi1Δ::HygR GFP-cnp1<sup>+</sup>:NatR sid4<sup>+</sup>-mRFP:KanR leu1-32 ura4-D18* | 3d |
| 7329 | *h+ pREP41-myc-his-ulp1<sup>+</sup> csi1Δ::HygR GFP-cnp1<sup>+</sup>:NatR sid4<sup>+</sup>-mRFP:KanR leu1-32 ura4-D18* | 3d |
| 7335 | *pREP41 csi1Δ::HygR nup132Δ::ura4<sup>+</sup> GFP-cnp1<sup>+</sup>:NatR sid4<sup>+</sup>-mRFP:KanR leu1-32 ura4-D18* | 3d |
| 7339 | *pREP41-myc-his-mature-pmt3<sup>+</sup> csi1Δ::HygR nup132Δ::ura4<sup>+</sup> GFP-cnp1<sup>+</sup>:NatR sid4<sup>+</sup>-mRFP:KanR leu1-32 ura4-D18* | 3d |
| 7341 | *pREP41-myc-his-mature-pmt3<sup>KallR</sup> csi1Δ::HygR nup132Δ::ura4<sup>+</sup> GFP-cnp1<sup>+</sup>:NatR sid4<sup>+</sup>-mRFP:KanR leu1-32 ura4-D18* | 3d |
| 7337 | *pREP41-myc-his-ulp1<sup>+</sup> csi1Δ::HygR nup132Δ::ura4<sup>+</sup> GFP-cnp1<sup>+</sup>:NatR sid4<sup>+</sup>-mRFP:KanR leu1-32 ura4-D18* | 3d |
| 8655 | *lem2<sup>+</sup>-GFP:KanR sid4<sup>+</sup>-mRFP:KanR leu1-32 ura4-D18/DSE* | 4a, 4b, S3b |
| 7689 | *h- nup132Δ::HygR lem2<sup>+</sup>-GFP:KanR sid4<sup>+</sup>-mRFP:KanR leu1-32 ura4-D18/DSE* | 4a, 4b, S3b |
| 7691 | *h+ csi1Δ::HygR lem2<sup>+</sup>-GFP:KanR sid4<sup>+</sup>-mRFP:KanR leu1-32 ura4-D18/DSE* | 4a, 4b |
| 7693 | *nup132Δ::HygR csi1Δ::HygR lem2<sup>+</sup>-GFP:KanR sid4<sup>+</sup>-mRFP:KanR leu1-32 ura4-D18/DSE* | 4a, 4b |
| 8639 | *lem2<sup>K7R</sup>-GFP:KanR sid4<sup>+</sup>-mRFP:KanR leu1-32 ura4-D18/DSE* | 4b, S3b |
| 7695 | *h- nup132Δ::HygR lem2<sup>K7R</sup>-GFP:KanR sid4<sup>+</sup>-mRFP:KanR leu1-32 ura4-D18/DSE* | 4b, S3b |
| 7744 | *csi1Δ::HygR lem2<sup>K7R</sup>-GFP:KanR sid4<sup>+</sup>-mRFP:KanR leu1-32 ura4-D18/DSE* | 4b |
| 7743 | *nup132Δ::HygR csi1Δ::HygR lem2<sup>K7R</sup>-GFP:KanR sid4<sup>+</sup>-mRFP:KanR leu1-32 ura4-D18/DSE* | 4b |
| 8717 | *bqt4Δ::NatR GFP-cnp1<sup>+</sup>:NatR sid4<sup>+</sup>-mRFP:KanR ura4-D18/DSE* | 4c |
| 8714 | *csi1Δ::HygR bqt4Δ::NatR GFP-cnp1<sup>+</sup>:NatR sid4<sup>+</sup>-mRFP:KanR ura4-D18/DSE* | 4c |
| 4 | *h- wild type S. pombe* | S1 |
| 5919 | *h+ nup132Δ::HygR lem2<sup>+</sup>-GFP:KanR ade6-210 leu1-32 ura4-D18/DSE imr1L(NcoI):ura4<sup>+</sup>* | S3a |
| 5908 | *h+ nup132Δ::HygR lem2<sup>K7R</sup>-GFP:KanR leu1-32 ura4-D18/DSE imr1L(NcoI):ura4<sup>+</sup>* | S3a |
| 5882 | *h+ lem2<sup>+</sup>-GFP:KanR leu1-32 ura4DS/E ade6-210 his1-102 imr1L(NcoI):ura4<sup>+</sup>* | S3a |
| 5884 | *h- lem2<sup>K7R</sup>-GFP:KanR leu1-32 ura4-D18/DSE imr1L(NcoI):ura4<sup>+</sup>* | S3a |
| 7679 | *slx8Δ::KanR nup132Δ::HygR GFP-cnp1<sup>+</sup>:NatR sid4<sup>+</sup>-mRFP:KanR leu1-32 ura4-D18/DSE* | S4 |
| 8000 | *slx8Δ::KanR csi1Δ::HygR sid4<sup>+</sup>-mRFP:KanR GFP-cnp1<sup>+</sup>:NatR ura4-D18 leu1-32* | S4 |
| 7745 | *csi1Δ::HygR nup132Δ::ura4<sup>+</sup> slx8Δ::KanR GFP-cnp1<sup>+</sup>:NatR sid4<sup>+</sup>-mRFP:KanR leu1-32 ura4-D18/DSE* | S4 |
| 8282 | *h+ ufd1ΔCt<sup>213-342</sup>:HygR nup132Δ::ura4<sup>+</sup> sid4<sup>+</sup>-mRFP:KanR GFP-cnp1<sup>+</sup>:NatR leu1-32 ura4-D18/DSE* | S4 |
| 8208 | *h- csi1Δ::KanR ufd1ΔCt<sup>213-342</sup>:HygR sid4<sup>+</sup>-mRFP:KanR GFP-cnp1<sup>+</sup>:NatR leu1-32 ura4-D18/DSE* | S4 |
| 8291 | *ufd1ΔCt<sup>213-342</sup>:HygR csi1Δ::KanR nup132Δ::ura4<sup>+</sup> sid4<sup>+</sup>-mRFP:KanR GFP-cnp1<sup>+</sup>:NatR leu1-32 ura4-D18/DSE* | S4 |

**Supplementary Table S3.** Plasmids used in this study.

| **Plasmid** | **Source** | **Figure** |
| --- | --- | --- |
| *pREP41-myc-his* | <sup>1</sup>Craven et al., 1998 | 2a, 2b, 3d |
| *pREP41-myc-his-ulp1<sup>+</sup>* | This study | 2a, 3d |
| *pREP41-myc-his-mature-pmt3<sup>+</sup>* | This study | 2b, 3d |
| *pREP41-myc-his-mature-pmt3<sup>KallR</sup>* | This study | 2b, 3d |

<sup>1</sup>Craven, R.A., D.J. Griffiths, K.S. Sheldrick, R.E. Randall, I.M. Hagan, and A.M. Carr. 1998. Vectors for the expression of tagged proteins in *Schizosaccharomyces pombe*. *Gene*. 221:59-68.

